# Structure-Activity Mapping of Intraperitoneal mRNA-LNPs: Decoupling Tumor and Liver Biodistribution in Pancreatic Cancer

**DOI:** 10.64898/2026.03.20.712457

**Authors:** Farhana Islam, Ashish Das, Md Ashaduzzaman, Ling Ding, Neha Kumari, Ran Dai, David Oupický

## Abstract

Pancreatic ductal adenocarcinoma (PDAC) remains difficult to treat with nucleic acid therapeutics because efficient intratumoral delivery is limited and off-target liver accumulation is common. Here, we developed a structure-activity map for intraperitoneally administered mRNA lipid nanoparticles (mRNA-LNPs) to identify formulation features that improve delivery to pancreatic tumors while reducing liver expression. A full-factorial library of 48 mRNA-LNP formulations was generated by varying ionizable lipid, sterol, phospholipid, and PEG-lipid components. Formulations were characterized for size, polydispersity, zeta potential, and encapsulation, then evaluated in an orthotopic KPC8060 pancreatic tumor model after intraperitoneal administration of firefly luciferase mRNA-loaded LNPs. Biodistribution was assessed by Rhodamine B fluorescence and functional delivery by luciferase expression 12 h after dosing.

Lipid composition strongly influenced both physicochemical properties and in vivo performance. G0-C14-based formulations produced the smallest and most homogeneous particles, whereas FTT5-containing formulations were generally larger. Across the 48-formulation library, mRNA expression and nanoparticle biodistribution varied significantly among tumor, pancreas, liver, and spleen. Statistical, decision-tree, and predictive modeling analyses identified composition rules associated with organ-selective delivery. High tumor expression was associated primarily with G0-C14 combined with DSPC and β-sitosterol, whereas liver expression was favored by C12-200 or DLin-MC3-DMA with DOPE and DSPE-PEG. Notably, a G0-C14/DSPC/DSPE-PEG formulation emerged as a lead candidate, producing a greater than 6-fold increase in tumor luciferase signal relative to the library median while reducing liver exposure by approximately 60%. Histopathology showed no treatment-related liver or lung toxicity. These findings define actionable formulation rules for tuning intraperitoneal mRNA-LNP delivery in PDAC and support further development of tumor-selective mRNA therapeutics for pancreatic cancer.

## INTRODUCTION

Lipid nanoparticles (LNPs) have transformed messenger RNA (mRNA) delivery for vaccines and gene therapies, but their propensity to accumulate in the liver after intravenous administration has driven the development of diverse strategies both with and without Selective Organ Targeting (SORT) to achieve precise delivery to organs like the pancreas, lungs, colon, heart, and brain. The SORT approach, which uses a specialized molecule to redirect LNPs, has enabled tailored targeting: for instance, lung-specific SORT LNPs with cationic lipids like DOTAP (50 mol%) bind vitronectin to deliver mRNA to lung cells, treating pulmonary lymphangioleiomyomatosis by restoring Tsc2 gene function in preclinical models (1), with protein expression peaking within hours. Similarly, colon targeting with SORT LNPs employs anionic lipids like 18PA to bind β2-glycoprotein I, directing mRNA to the colon for potential colorectal cancer or inflammatory bowel disease therapies, evidenced by enhanced Cy5-mRNA signals in preclinical studies (2). In contrast, non-SORT methods leverage alternative lipid compositions and administration routes demonstrated pancreatic targeting without SORT (3), using LNPs with ionizable lipids (e.g., 306O) and cationic helper lipids like DOTAP, delivered intraperitoneally to C57BL/6 mice, achieving up to 60% specificity in pancreatic β cells via macrophage-mediated exosome transfer, peaking at 3-4 hours, with applications for diabetes or pancreatic cancer. Beyond SORT, heart targeting has been achieved through intravenous LNPs with optimized ionizable lipids like MC3 (4) delivering mRNA to cardiomyocytes in mice for cardiac repair, bypassing the need for a SORT molecule by fine-tuning lipid ratios (e.g., 50:10:38.5:1.5 MC3:DSPC:cholesterol:PEG-DMG). Likewise, brain targeting without SORT uses neutral lipids like DOPE in LNPs, delivered intranasally or via focused ultrasound to cross the blood-brain barrier achieving mRNA expression in neurons for neurological disorders like Alzheimer’s (5). Liver targeting, a default for many LNPs, is enhanced by ApoE binding in both SORT and non-SORT contexts, as with patisiran’s FDA-approved siRNA delivery (6). Clinical trials like BioNTech’s BNT111 for melanoma further validate LNP potential, while SORT and non-SORT strategies together delivering multiple mRNAs (e.g., luciferase, GFP, mCherry) or single payloads offer versatile, low-toxicity options to revolutionize organ-specific therapies, from respiratory and gastrointestinal diseases to cardiac and neurological conditions.

Pancreatic ductal adenocarcinoma (PDAC) is the predominant histological subtype of pancreatic cancer (PC), comprising 90% of cases and carrying a 5-year survival rate of only 12% (8, 9). By 2030, PDAC is projected to become the second-leading cause of cancer-related mortality in the United States (10). In 2023, 64,050 new cases were estimated in the U.S., with 50,550 deaths attributed to the disease (11). The lack of specific early symptoms and the aggressive nature of PDAC contribute to delayed diagnoses, limited therapeutic responses, and poor outcomes (12, 13). The fibrotic tumor microenvironment (TME) serves as a significant barrier to drug delivery, reducing the efficacy of standard chemotherapeutic agents (14). Additionally, the TME’s immunosuppressive nature limits the success of immunotherapies, such as checkpoint inhibitors and CAR T cells, which have been effective in other solid tumors but show minimal efficacy in PDAC (15). Surgery remains the only potentially curative option; however, less than 20% of patients qualify due to the advanced disease stage at presentation. Chemotherapy, often administered to reduce relapse, is limited by its severe side effects and uncertain ability to overcome TME-associated barriers (10). Recent breakthroughs in immunotherapy and novel drug delivery systems provide promising direction for PDAC treatment, emphasizing the need for innovative approaches to improve therapeutic efficacy and patient outcomes (16).

mRNA has emerged as a promising therapeutic modality. Unlike DNA-based therapies, mRNA does not integrate into the host genome, reducing the risk of insertional mutagenesis (17, 18). Its transient nature allows for controlled protein expression, and the cell-free production process enables rapid and scalable manufacturing. These attributes make mRNA a versatile platform for developing personalized cancer vaccines and immunotherapies (19). In oncology, mRNA-based therapies offer several benefits. They can be tailored to encode tumor-specific antigens or therapeutic proteins directly to cancer cells, enhancing targeting accuracy and minimizing off-target effects, while combining immune modulation with tumor-specific treatment to improve therapeutic efficacy and reduce systemic toxicity. However, challenges remain in the clinical application of mRNA therapeutics. The inherent instability of mRNA makes it susceptible to degradation, necessitating the development of effective delivery systems to protect the mRNA and facilitate its uptake by target cells (21, 22). The delivery of mRNA therapeutics to PDAC remains particularly challenging due to the tumor’s dense fibrotic stroma, which acts as a physical and biochemical barrier (23). The excessive deposition of extracellular matrix components, including collagen and hyaluronan, increases interstitial fluid pressure and limits nanoparticle penetration (24). Additionally, PDAC is characterized by a highly immunosuppressive TME that further impairs therapeutic efficacy (25). LNPs have shown promise in protecting mRNA from enzymatic degradation and facilitating cellular uptake. Strategies such as hyaluronidase-based stromal depletion, receptor-targeted nanoparticles (e.g., integrin or fibroblast activation protein-targeted LNPs), and pH-responsive carriers are being explored to enhance mRNA delivery, improve endosomal escape, and increase therapeutic efficacy in PDAC (26).

LNPs have been employed to address the delivery issues and optimizing these delivery vehicles for efficient and safe administration and tissue-specific delivery continues to be an area of active research (27). Furthermore, while mRNA-based therapies, including vaccines, have shown promise in clinical trials, ensuring consistent and sustained therapeutic responses across diverse patient populations remains a significant challenge, with successful integration into standard oncology practice requiring the overcoming of these hurdles (20). Advancements in mRNA-based nanotechnology have revolutionized cancer therapeutics, offering innovative strategies to combat tumors. mRNA-encoded tumor suppressors halt cancer cell proliferation (28, 29), while mRNA-encoded antigens and cytokines activate immune responses within the host (30–32). Additionally, mRNA-encoded genome editing proteins target tumor survival genes, enhancing treatment efficacy with fewer interventions. Promising results have also been observed with mRNA-encoded CARs and TCRs for T cell engineering in cancer therapy (33). Despite these breakthroughs, challenges such as mRNA’s inherent instability, immunogenicity, and delivery difficulties have historically limited its progress (34). LNPs have emerged as a promising solution to these obstacles. By encapsulating mRNA, LNPs protect it from degradation and facilitate its entry into target cells (35–37). Recent advancements have enhanced LNP design to improve organ-specific delivery. For instance, researchers have developed LNPs capable of crossing the blood-brain barrier, enabling targeted mRNA delivery to brain cells, a significant step toward treating neurological diseases like Alzheimer’s (38). Additionally, LNPs have been engineered to deliver mRNA therapeutics to the placenta, offering potential treatments for conditions such as pre-eclampsia (39).

We aim to develop a comprehensive library of LNP formulations, varying lipid composition to optimize the targeted delivery of mRNA therapeutics through intraperitoneal (IP) administration. This decision is driven by our previous findings that physicochemical attributes significantly impact the biodistribution of formulations (43) and IP delivery is more effective than intravenous (IV) administration when targeting pancreatic tumors (44). One study demonstrated that IP administration of LNPs with cationic helper lipids like DOTAP enables potent and specific mRNA delivery to pancreatic β cells via macrophage-mediated exosome transfer, offering a promising non-viral gene therapy approach for pancreatic diseases such as diabetes and cancer (3). Developing a comprehensive formulation library is essential for optimizing LNP compositions to achieve targeted delivery of mRNA therapeutics. For instance, modifying the lipid composition can alter the surface charge and hydrophobicity of LNPs, thereby enhancing their ability to target particular organs (45). This approach also facilitates the identification of optimal LNP formulations that maximize therapeutic efficacy while minimizing off-target effects (46). Understanding the relationships between LNP formulation parameters, biodistribution, and mRNA expression through correlation analysis enables to systematically identify key factors that enhance the efficiency and targeting of mRNA delivery systems (47, 48).

In this project, the aim is to advance the development of LNP formulations for mRNA delivery by achieving targeted delivery. Specifically, the project seeks to systematically assess the biodistribution and mRNA expression across various organs to determine how formulation parameters influence delivery outcomes. Correlation analysis will be performed to identify key relationships between biodistribution, mRNA expression, and LNP composition, providing insights into structure-activity relationships. Through experimental and statistical analysis, the project aims to identify optimal LNP compositions that balance stability, cellular uptake, and precise targeting. The project is outlined in the pictorial study overview illustrated in (Fig. 1). These findings will refine LNP designs for maximal efficacy in specific therapeutic contexts, such as oncology or infectious diseases.

**Fig. 1.**
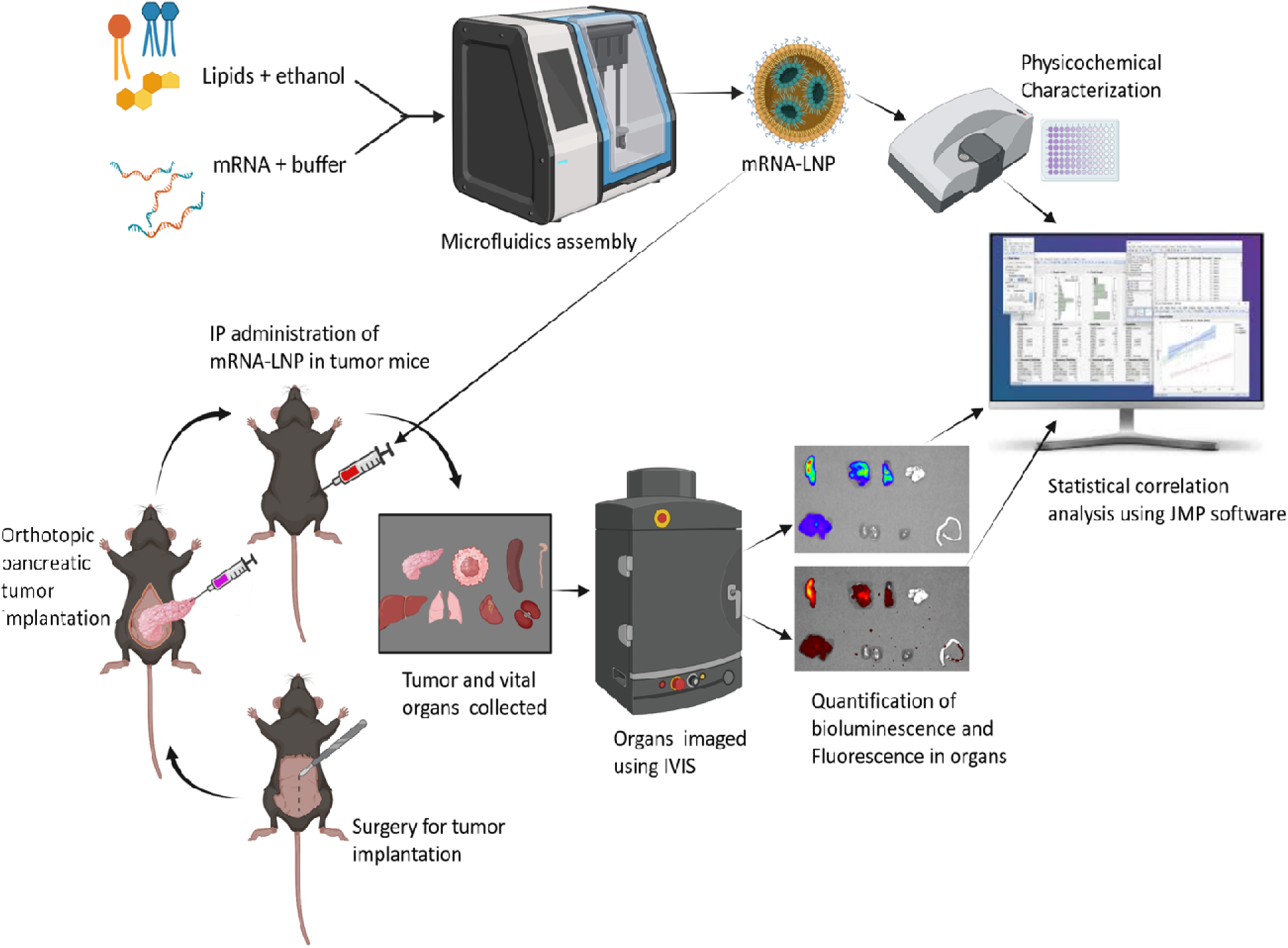
Schematic overview of the experimental workflow for mRNA-LNP formulation, characterization, and *in vivo* biodistribution and mRNA expression evaluation. Lipid nanoparticles (LNPs) were prepared by mixing lipids dissolved in ethanol with mRNA in an aqueous buffer using a microfluidic assembly system. The resulting mRNA-LNP formulations were characterized for size, polydispersity index (PDI), zeta potential (via dynamic light scattering), and encapsulation efficiency (via RiboGreen assay). For in vivo studies, orthotopic pancreatic tumor models were established in mice through surgical implantation of tumor tissue. mRNA-LNPs were administered intraperitoneally (IP) into tumor-bearing mice. After treatment, tumor and vital organs were collected, and fluorescence and bioluminescence were quantified using an in vivo imaging system (IVIS). The collected data were further analyzed using JMP software to evaluate statistical correlations between mRNA expression and biodistribution. This integrated approach facilitated a comprehensive assessment of the physicochemical properties and in vivo performance of the mRNA-LNP formulations.

## MATERIALS AND METHODS

### Materials

[(6Z,9Z,28Z,31Z)-heptatriaconta-6,9,28,31-tetraen-19-yl] 4-(dimethylamino)butanoate (DLinMC3) was purchased from Cayman Chemical (Ann Arbor, Michigan, USA, Item# 34364). 1,1-((2-(4-(2-((2-(bis(2-hydroxydodecyl)amino)ethyl)(2-hydroxydodecyl) amino)ethyl)piperazin-1-yl)ethyl)azanediyl)bis(dodecan-2-ol) (C12-200) was obtained from MedKoo Biosciences, Inc. (Durham, NC, USA, Cat#: 556048). Octan-4-yl 9-[3-[[3,5-bis[3-[bis(9-octan-4-yloxy-9-oxononyl)amino]propylcarbamoyl]benzoyl]amino]propyl-(9-octan-4-yloxy-9-oxononyl)amino]nonanoate (FTT5) and 3-[[3-[2-[bis(2-hydroxytetradecyl)amino]ethylamino]-3-oxopropyl]-[2-[[3-[2-[bis(2-hydroxytetradecyl)amino]ethylamino]-3-oxopropyl]-[3-[2-(2-hydroxytetradecylamino)ethylamino]-3-oxopropyl]amino]ethyl]amino]-N-[2-(2-hydroxytetradecylamino)ethyl]propanamide (G0-C14) were purchased from MedChemExpress LLC (Monmouth Junction, NJ, USA, Cat# HY-145793 and HY-152229, respectively). Cholesterol was obtained from Thermofisher Scientific (Waltham, MA, Cat# A11470.0B), and β-sitosterol was purchased from Sigma-Aldrich (St. Louis, MO, Cat# S1270, CAS# 83-46-5). 20α-Hydroxycholesterol was acquired from MedChemExpress LLC (Monmouth Junction, NJ, USA, Cat# HY-12316). Distearoyl-phosphatidylethanolaminemethyl-polyethyleneglycol conjugate 2000 (DSPE-PEG2000) and [3-(2-methoxyethoxy)-2-tetradecanoyloxypropyl] tetradecanoate (DMG-PEG2000) were obtained from NOF Corporation (Shibuya-ku, Tokyo, Japan, Cat# DSPE-020CN and GM-020, respectively). Distearoylphosphatidylcholine (DSPC) was purchased from Avanti Polar Lipids (Alabaster, AL, USA, Cat# 850365), and 1,2-Dioleoyl-sn-glycero-3-phosphoethanolamine (DOPE) was sourced from NOF Corporation (Shibuya-ku, Tokyo, Japan, Cat# ME-8181). Rhodamine B 1,2-Dihexadecanoyl-sn-Glycero-3-Phosphoethanolamine, Triethyl-ammonium Salt (Rhodamine B-DHPE) was purchased from Thermofisher Scientific (Waltham, MA, Cat# L1392). Pure ethanol was used in the study. Firefly Luciferase mRNA (fLuc mRNA) and Enhanced Green Fluorescent Protein mRNA (EGFP mRNA) were obtained from TriLink Biotechnologies (San Diego, California, USA, Cat# L-7602 and L-7601, respectively). 100 mM pH 4 citrate buffer was sourced from MedChemExpress LLC (Monmouth Junction, NJ, USA, Cat# HY-B1610N). A 10 kDa MWCO Amicon® Ultra Centrifugal Filter (Millipore Sigma, Cat# UCF901024) was used, along with 96% paraformaldehyde from Thermofisher Scientific (Waltham, MA, Cat# 416780100). The Ribogreen reagent kit was obtained from Invitrogen (Thermofisher Scientific, Waltham, MA, Cat# R11490). D-Luciferin, Messengermax lipofectamine kit, Dulbecco’s Modification of Eagle’s Medium (DMEM) cell culture media, Fetal Bovine Serum (FBS), Penstrep, Trypsin, Opti-MEM, and Matrigel matrix were all purchased from Thermofisher Scientific (Waltham, MA, Cat# LMRNA008, 31985062, FB12999102, SH30042.01, and CB-40234A, respectively) and Cytiva/HyClone (Marlborough, MA, USA, Cat# SH30256.01). Phosphate-buffered saline (PBS) was also purchased from Cytiva/HyClone (Marlborough, MA, USA, Cat# SH30256.01). Hematoxylin (Fisher Scientific, Cat# STLSL401) and Eosin (Fisher Scientific, Cat# STLSL201), Buprenorphine ER 1mg/ml (Veterinarian prescribed), Isoflurane, USP (Piramal Critical Care, Inc., Bethlehem, PA, USA, NDC 66794-013-25).

## Methods

### Formulation design and LNP preparation

A full-factorial 4 × 3 × 2 × 2 design (ionizable lipid × sterol × phospholipid × PEG-lipid) generated 48 unique formulations. For every formulation the molar ratio of ionizable : sterol : phospholipid : PEG-lipid : fluorescent tracer (RhB-DHPE) was fixed at 50 : 37.5 : 10 : 1.5 : 1. mRNA-loaded lipid nanoparticles (mRNA-LNPs) were formulated using a microfluidics assembly method with NanoAssemblr™ Ignite™ (Cytiva, Marlborough, MA, USA). Briefly, 150 µg of mRNA was dissolved in a 600 µl 100 mM citrate buffer at pH 4.0. The lipid components including cationic ionizable lipids (C12-200/ DLin-MC3/ FTT5/ G0-C14), helper lipids (cholesterol/β-sitosterol/20α-hydroxycholesterol), phospholipids (DOPE/ DSPC), PEGylated lipids (DMG-PEG2000/ DSPE-PEG2000), and Rhodamine B-DHPE were dissolved in pure ethanol at a molar ratio 50:37.5:10:1.5:1. The physicochemical properties such as M.W, Log*P* values and chemical structure of these lipids are showed in (Table 1) (52–61) . The organic and aqueous phases were then introduced into a microfluidics chip at a flow rate ratio (FRR) of 3:1, with a total flow rate (TFR) of 12 mL/min. Following the assembly, the mRNA-LNP formulation was diluted 40-fold with phosphate-buffered saline (PBS) and subsequently concentrated by centrifugation using a 10 kDa MWCO Amicon ultra centrifugal filter and centrifuge instrument Sorvall Legend XTR (Thermo Fisher Scientific Inc., Waltham, MA). We made a library of 48 mRNA-LNP formulations by varying lipid composition within the LNP formulations (Table S1).

**Table 1.**
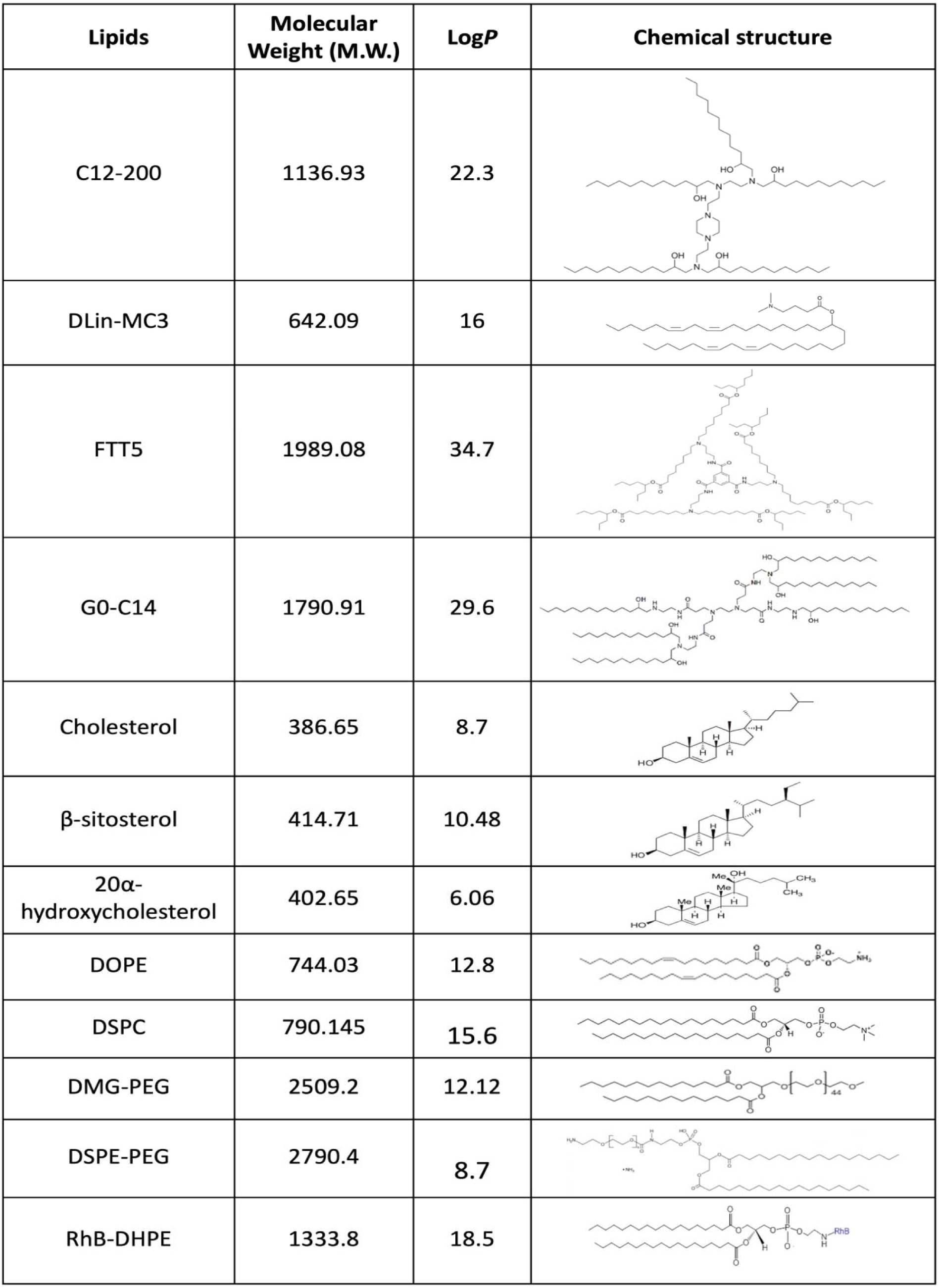
Physicochemical properties of lipids used in the mRNA-LNP library.

| Lipids | Molecular Weight (M.W.) | LogP | Chemical structure |
| --- | --- | --- | --- |
| C12-200 | 1136.93 | 22.3 |  |
| DLin-MC3 | 642.09 | 16 |  |
| FTT5 | 1989.08 | 34.7 |  |
| G0-C14 | 1790.91 | 29.6 |  |
| Cholesterol | 386.65 | 8.7 |  |
| β-sitosterol | 414.71 | 10.48 | |
| 20α-hydroxycholesterol | 402.65 | 6.06 | |
| DOPE | 744.03 | 12.8 |  |
| DSPC | 790.145 | 15.6 |  |
| DMG-PEG | 2509.2 | 12.12 |  |
| DSPE-PEG | 2790.4 | 8.7 |  |
| RhB-DHPE | 1333.8 | 18.5 |  |

### Physicochemical characterization

Particle size, polydispersity index (PDI), and zeta potential were determined using dynamic light scattering (DLS) with a Zeta Sizer Nano ZS (Malvern Nano-ZS, Worcestershire, UK). Each formulation was diluted 10-fold in 1x PBS to achieve optimal scattering intensity, and three readings of the same sample were recorded. Encapsulation efficiency (EE%) of mRNA in the LNP formulations was quantified using the RiboGreen® RNA Assay (Invitrogen, Thermofisher Scientific, Waltham, MA, USA). A standard curve of known mRNA concentrations was prepared for accurate quantification of encapsulated versus free mRNA. Formulations were treated with 1% Triton X-100 to lyse the particles and release encapsulated mRNA, followed by fluorescence measurement at excitation and emission wavelengths of 485 nm and 530 nm, respectively, using a microplate reader SpectraMax iD3 (Molecular Devices, LLC., San Jose, CA, 95134, USA). The encapsulation efficiency was calculated as the ratio of encapsulated mRNA to the total mRNA content.

### Cells

The mouse PC cell line KPC8060 was generously provided by Dr. Hollingsworth at the University of Nebraska Medical Center (UNMC). The cells were maintained at 37 °C with 5% CO in a humidified incubator, using high-glucose DMEM supplemented with 10% FBS and penicillin-streptomycin (100 U/mL, 100 μg/mL). For subculturing, cells were detached with trypsin, then centrifuged using Sorvall Legend XTR (Thermo Fisher Scientific Inc., Waltham, MA, USA) at 1000 rpm for 5 mins to pellet the cells.

### Animal studies

Male C57BL/6 mice (7 weeks old, 20g body weight) were obtained from Charles River Laboratories. The animals were housed in the UNMC Comparative Medicine facilities with access to food and water. All animal experiments were conducted with the approval of the UNMC Institutional Animal Care and Use Committee (IACUC). We adhered to all applicable federal ethical regulations for the use of animals in research, ensuring the ethical treatment and care of laboratory animals in accordance with protocol 18-045-05. These animals were used to conduct the mRNA-LNP biodistribution and mRNA expression analysis.

Orthotopic tumors were generated in C57BL/6 mice using KPC8060 cells. The mice were anesthetized with isoflurane D throughout the surgical procedure. The surgical site was sterilized with iodine and alcohol pads. A midline incision was made in the left central abdominal region to access the pancreas. A mixture of 40 μL Matrigel/PBS (1:1) containing 2.5 × 10 KPC8060 cells was injected into the pancreatic tail. The incision was closed in two layers: the inner layer with 5−0 chromic catgut sutures and the outer layer with soft staples. Buprenorphine ER was administered subcutaneously at a dose of 1mg/kg for postoperative pain management. 8 days after surgery, the soft staples were removed.

Tumor progression was monitored using high-resolution ultrasound imaging on alternate days, beginning on day 8 post-implantation. By day 13, when the pancreatic tumors reached an approximate diameter of 1 cm, as confirmed by ultrasound, the mice were treated with an intraperitoneal injection of the mRNA-LNP formulation. The formulation was administered at a dose of 0.5 mg/kg mRNA using U-100 insulin syringes fitted with 28G x 1/2” needles to ensure accurate and minimally invasive delivery. 12 hours post-treatment, the mice were euthanized following institutional ethical guidelines for animal studies. Pancreatic tumors, along with major organs including the liver, lungs, healthy pancreas, spleen, colon, heart, and kidneys, were carefully excised and processed for downstream imaging. This approach facilitated the assessment of toxicity profiling, biodistribution and mRNA expression.

The mRNA-LNP formulations were designed for comprehensive biodistribution and functional assessment. To enable tracking, the formulations were labeled with Rhodamine B (RhB), a fluorescent dye, which facilitated visualization of their distribution in tissues. Additionally, the formulations were loaded with fLuc mRNA to evaluate the expression of mRNA in targeted and non-targeted tissues. Control mice were treated with phosphate-buffered saline (PBS) as a negative control to establish baseline fluorescence and bioluminescence levels. 12 hours after treatment, the tumor-bearing mice were euthanized following institutional animal care and use guidelines. Organs, including the pancreatic tumor, liver, lungs, healthy pancreas, spleen, colon, heart, and kidneys, were carefully harvested, and processed for imaging. Special attention was given to preserving the integrity of each organ to ensure accurate and reliable imaging results.

The collected organs were imaged ex vivo using the IVIS® Spectrum in vivo imaging system (PerkinElmer, Waltham, MA, USA), a highly sensitive tool capable of capturing both fluorescence and bioluminescence signals. RhB fluorescence was detected at an excitation wavelength of 571 nm and an emission wavelength of 591 nm, while fLuc mRNA expression was measured through bioluminescence intensity. These dual modalities provided complementary data, with fluorescence indicating the biodistribution of the LNP formulations and bioluminescence revealing the functional expression of the delivered mRNA. Quantitative analyses of the fluorescence and bioluminescence signals were performed using Living Image® 4.5 software. This software enabled precise spatial mapping and intensity measurement of the signals within each organ, providing a detailed biodistribution profile and insights into the formulation’s efficiency in mRNA delivery and expression. The combined use of fluorescence and bioluminescence allowed for a robust assessment of both physical localization and biological activity of the mRNA-LNP formulations.

### *In vivo* toxicity assay

Tumors and major organs harvested from the tumor mice were preserved. Tissues were sectioned via microtome at 4 microns on to a positively charged slide, dried at room temp, and then baked at 60°C. Slide were rehydrated and stained with Hematoxylin to stain the cell nuclei and Eosin to stain the cytoplasm and extracellular matrix. Following staining, the slides were dehydrated, and cover slipped with Tissue-Tek Prima/Glas. Finally, the prepared slides were imaged by a pathologist and observations were listed.

### Statistical analysis

We investigated the pairwise association among the LNP formulations, mRNA expression, LNP biodistribution, particle size, PDI, and zeta potential. All statistical analyses were conducted using JMP software (JMP® Pro version 18.0.2, JMP Statistical Discovery LLC).

To assess the impact of different lipid components on LNP size, zeta potential, and PDI, we employed Kruskal-Wallis test and ANOVA analysis. Specifically, the Kruskal-Wallis test was employed to assess statistically significant differences in the median sizes across the different lipids within each lipid type. ANOVA analysis was performed to assess whether mean size differences exist among different lipids of the lipid types.

To identify composition-based classification rules, decision tree analysis was conducted using the Partition method in JMP. LNP size, zeta potential, and PDI were categorized into predefined ranges, and a pruned decision tree was generated to highlight the most significant lipid components influencing these properties.

A Support Vector Regression (SVR) model was developed to analyze the relationship between lipid composition and LNP size. Grid search optimization was performed to fine-tune hyperparameters such as cost and gamma, ensuring improved predictive accuracy. Interaction effects between lipid types were visualized through interaction profiler plots, providing insights into how lipid combinations influence nanoparticle characteristics. The same methodology was applied to model and predict zeta potential and PDI values.

To evaluate the impact of lipid composition on LNP biodistribution and mRNA expression across different organs, we conducted an ANOVA where fluorescence intensity (for biodistribution) and expression levels (for mRNA) were analyzed as response variables, with LNP formulation as the independent variable. A blocking factor was introduced to account for inter-mouse variability, ensuring consistency and reproducibility of results. Data was normalized across organs to standardize variations.

Biodistribution and mRNA expression data were categorized into high, medium, and low expression groups. A bar graph visualization was generated to illustrate the distribution of formulations achieving different expression levels across organs. To assess lipid-specific influences on biodistribution patterns, a Kruskal-Wallis test was applied to compare median fluorescence intensity across lipid groups, and an ANOVA was conducted to analyze mean differences.

A decision tree analysis was further implemented to classify formulations based on their biodistribution and mRNA expression profiles. Lipid composition served as the predictor variable, while fluorescence intensity and expression levels were classified into categorical response variables. The decision tree was pruned to retain the most significant rules, enabling the identification of key lipid components driving high or low biodistribution in specific organs.

To explore potential relationships among LNP size, PDI, zeta potential, biodistribution, and mRNA expression, we performed Pearson’s correlation analysis. The correlation coefficient (r) was calculated to assess the strength and direction of associations among these variables, with predefined thresholds for strong, moderate, and weak correlations. This analysis provided insights into how physicochemical properties of LNPs influence their biological behavior.

All statistical analyses were conducted with a significance threshold of p < 0.05. Detailed statistical results, effect sizes, and specific formulation trends are reported in the Results section.

## RESULTS

### mRNA-LNP synthesis and characterization

The hydrodynamic diameter and PDI provided insights into particle uniformity, while zeta potential, determined by electrophoretic light scattering, assessed surface charge and colloidal stability. The size distribution of LNP formulations, grouped by the cationic ionizable lipid used, displayed distinct trends (Fig. 2a-d). For C12-200-based formulations, nine out of twelve formulations exhibited particle sizes below 150 nm, highlighting its ability to consistently produce small nanoparticles (Fig. 2a). DLin-MC3-DMA-based formulations showed a broader size range, with six formulations measuring below 150 nm and six between 150-250 nm (Fig. 2b). In the case of FTT5-based formulations, most formulations were larger, with sizes ranging from 150-350 nm, suggesting challenges in achieving smaller particle sizes with this lipid (Fig. 2c). Conversely, G0-C14-based formulations demonstrated the smallest particle sizes, with ten out of twelve formulations measuring below 120 nm, showcasing their efficiency in forming small and uniform nanoparticles (Fig. 2d). These results indicate that the choice of cationic ionizable lipid significantly influences the size distribution of LNP formulations.

**Fig. 2.**
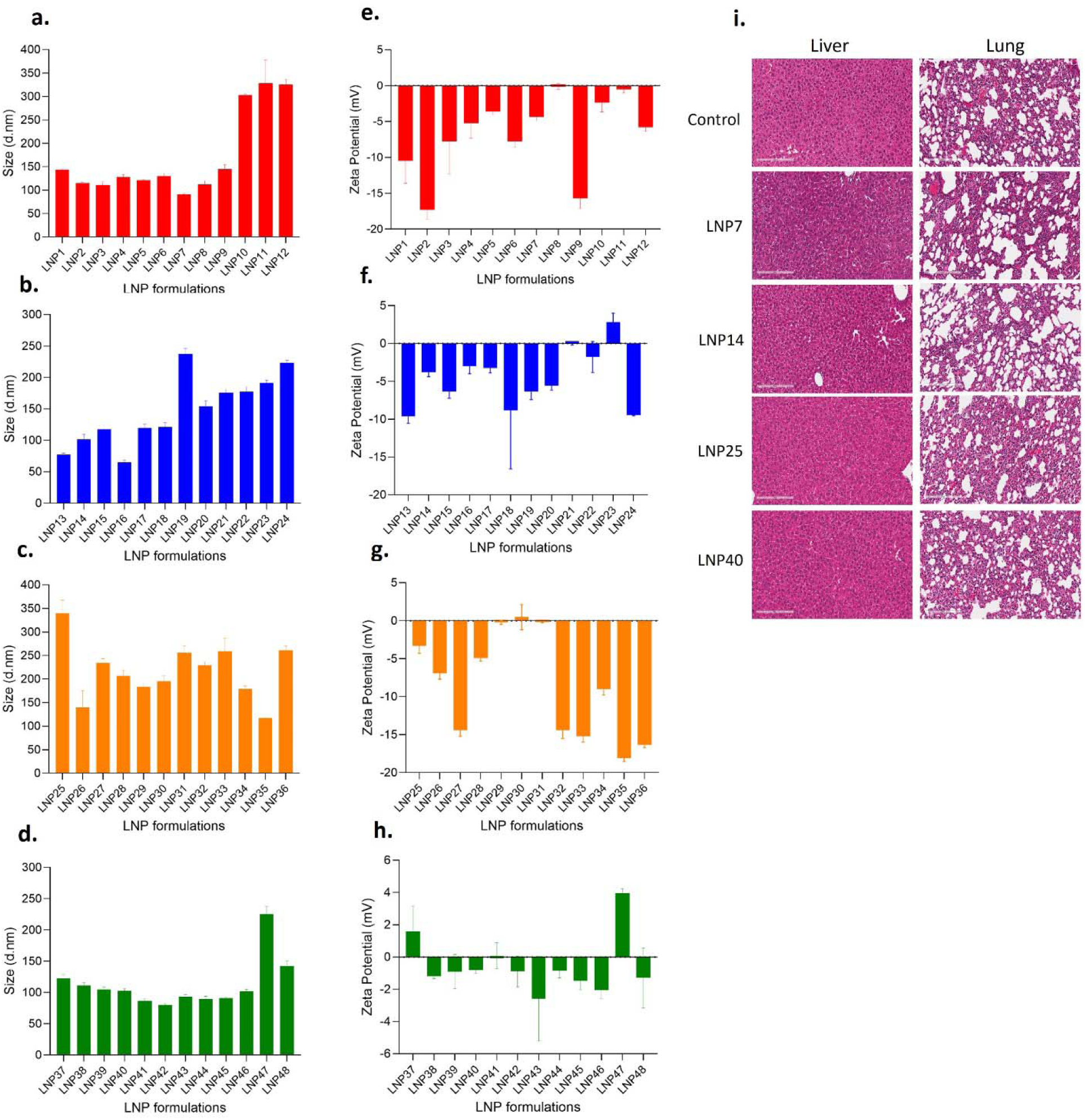
Characterization and toxicity assessment of 48 mRNA-LNP formulations grouped in four different cationic ionizable lipids. (a-d) Hydrodynamic size (d.nm) of mRNA-loaded lipid nanoparticles (LNPs) formulated with C12-200 (a, red), DLin-MC3-DMA (b, blue), FTT5 (c, orange), and G0-C14 (d, green), measured via dynamic light scattering (DLS). Each lipid library contains 12 formulations. (e-h) Zeta potential (mV) of mRNA-LNP formulations prepared with C12-200 (e, red), DLin-MC3-DMA (f, blue), FTT5 (g, orange), and G0-C14 (h, green), measured via electrophoretic mobility. (i) Hematoxylin and Eosin (H&E) stain images of liver (top panel) and lung (bottom panel) tissues from mice administered mRNA-LNP formulations. Lung and liver tissues from four different treatment groups (LNP7, LNP14, LNP25 and LNP40) were compared to tissues from untreated mice. The left panels show liver morphology, and right panels show lung morphology. Scale bar: 200 μm.

The zeta potential of the formulations was also evaluated to understand their surface charge characteristics, which influence stability and interaction with biological systems (Fig. 2e-h). C12-200 formulations showed a predominantly negative zeta potential, ensuring colloidal stability in suspension (Fig. 2e). DLin-MC3-DMA formulations exhibited varied zeta potentials, with most values near neutrality or slightly negative, suggesting moderate stability (Fig. 2f). FTT5 formulations demonstrated more negative zeta potentials, consistent with their larger size, possibly reflecting the lipid composition’s effects on surface charge (Fig. 2g). G0-C14-based LNPs consistently exhibited near neutral zeta potentials ranging from ∼ −5 to +3 mV, reinforcing the trend of higher colloidal stability seen with their smaller particle size (Fig. 2h).

The PDI of the 48 fLuc mRNA-LNP formulations was evaluated to assess the uniformity of particle size distribution across four categories of cationic ionizable lipids (Fig. S1a-d). Formulations prepared with C12-200 exhibited increasing PDI values across formulations, with some approaching 0.4, indicating moderate variability in particle size distribution (Fig. S1a). DLin-MC3-DMA formulations exhibited a wider range of PDI values, with some below 0.1, others between 0.1 and 0.2, and a few nearing 0.3, reflecting a more variable size distribution (Fig. S1b). FTT5-based formulations displayed relatively low PDI values, with most formulations below 0.2, though several exceeded 0.3, signifying considerable heterogeneity in particle size (Fig. S1c). Notably, G0-C14 (Fig. S1d) formulations achieved the most uniform particle size distribution, with PDI consistently below 0.2, highlighting its higher ability to produce highly homogeneous nanoparticles. These results demonstrate that the choice of cationic ionizable lipid plays a critical role in determining the size uniformity of LNP formulations, with G0-C14 showing the best performance in achieving high homogeneity.

The encapsulation efficiency (EE%) of the 48 fLuc mRNA-LNP formulations was systematically evaluated across four groups of cationic ionizable lipids, revealing high levels of encapsulation across all formulations (Fig. S1e-h). C12-200-based formulations exhibited EE% values ranging from 82.4% to 98%, with the majority maintaining values above 85%, demonstrating consistent encapsulation efficiency (Fig. S1e). Similarly, DLin-MC3-DMA-based formulations achieved EE% values between 83% and 98%, with most formulations exceeding 90%, indicating robust performance (Fig. S1f). FTT5-based formulations showed EE% values ranging from 81% to 98%, with most formulations above 85%, though with slightly greater variability (Fig. S1g).

Despite this variability, these three groups consistently achieved effective mRNA encapsulation. In contrast, G0-C14-based formulations demonstrated the highest level of consistency, with EE% values ranging from 89% to 98%, underscoring its superior ability to maintain uniform and high encapsulation efficiency (Fig. S1h).

### *In vivo* toxicity profiling

The toxicity of mRNA-LNP formulations was evaluated in male C57 BL/6 mice with orthotopically implanted pancreatic tumors. Liver and lung tissue sections were collected 12 hours post-treatment and stained with hematoxylin and eosin (H&E). A pathologist conducted a comprehensive histopathological evaluation to assess treatment-related toxicity (Table S6).

In the control group (untreated mice), pathological findings were observed in the liver. These included “tumor in liver and on capsule” and “small tumor on capsule,” which are consistent with the progression of PC and the expected metastatic or local tumor spread in this model. In contrast, all lung tissue sections in the control group appeared normal, with no evidence of metastatic lesions or other abnormalities, indicating that the disease progression had not significantly affected the lungs by the time of analysis. In the treatment groups (mRNA-LNP formulations), liver histology revealed that most treated animals displayed normal tissue architecture, with no evidence of hepatotoxicity. This suggests that the formulations did not induce liver toxicity within the study timeframe. A few liver samples showed findings such as “tumor in liver and on capsule,” consistent with the natural disease progression and not indicative of treatment-related toxicity, as similar findings were present in the control group. Lung tissue in the treatment groups exhibited normal histology, with no signs of toxicity such as inflammation, necrosis, or fibrosis. This demonstrates the absence of pulmonary toxicity following mRNA-LNP administration.

In summary, the liver and lung histology in the treatment groups closely mirrored that of the control group, with no additional pathological changes attributable to the LNP formulations (Fig. 2i). The presence of liver tumors in both groups reflect the expected progression of the pancreatic tumor model and does not signify treatment-induced toxicity. Lung tissues in both treated and untreated groups remained unaffected, reinforcing the safety profile of the tested LNP formulations. This histopathological analysis confirms that mRNA-LNP formulations are well-tolerated. No treatment-related toxicity was observed in either liver or lung tissues when compared to controls. Observed liver abnormalities, such as tumor presence, are attributed to the inherent characteristics of the orthotopic pancreatic tumor model rather than the treatments. These findings support the safety of the formulations and their suitability for further preclinical evaluation.

### Biodistribution and mRNA expression analysis

The biodistribution and mRNA expression of fLuc mRNA-LNPs were evaluated in male C57BL/6 mice with orthotopically implanted pancreatic tumors using KPC8060 cells. Following tumor growth to approximately 1 cm, monitored via ultrasound, mice received intraperitoneal injections of LNPs loaded with 0.5 mg/kg of mRNA. Analysis was performed 12 hours post-administration using the IVIS imaging system (Fig. 3a).

**Fig. 3.**
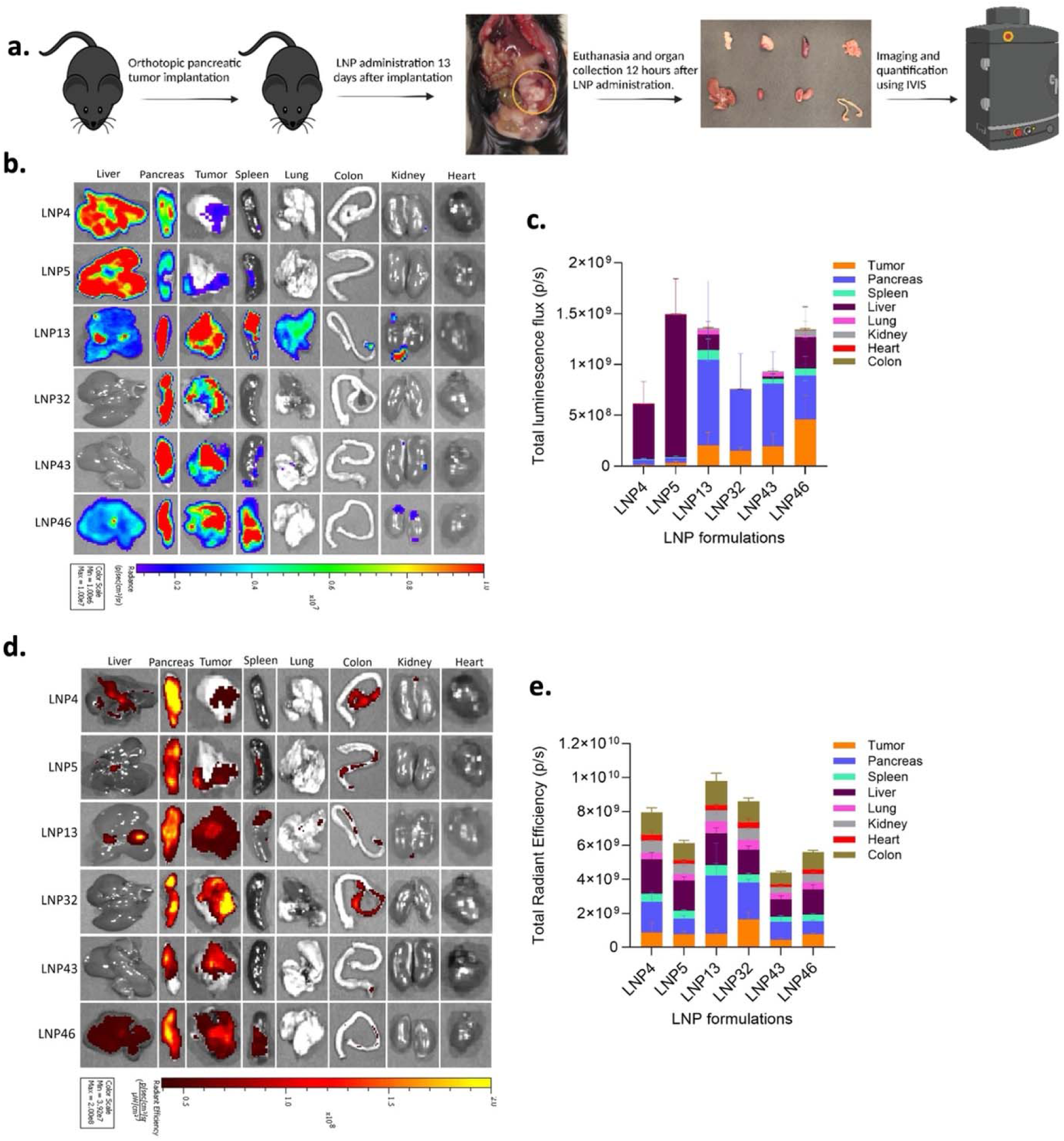
Biodistribution and mRNA expression analysis of Fluc mRNA-loaded LNPs in orthotopic pancreatic tumor-bearing mice. (a) Schematic representation of the experimental workflow. Orthotopic pancreatic tumors were implanted in male C57BL/6 mice using KPC8060 cells. Tumor growth was monitored via ultrasound every alternate day. When tumors reached ∼1 cm, mice were intraperitoneally administered Fluc mRNA-loaded LNPs (0.5 mg/kg mRNA). Mice were euthanized 12 hours post-administration, and organs, including tumors, were collected for imaging and analysis using the IVIS instrument. (b) Representative bioluminescence images showing Fluc mRNA expression across major organs including liver, pancreas, spleen, lung, colon, kidney, and heart and tumors for selected LNP formulations (LNP4, LNP5, LNP13, LNP32, LNP43, LNP46). (c) Quantification of Fluc mRNA expression (total luminescence flux, p/s) in major organs and tumors using Living Image software. Data are represented as mean ± SD (n=5). (d) Representative images showing RhB fluorescence (LNP biodistribution) across major organs and tumors for the same formulations. (e) Quantification of RhB fluorescence intensity (total radiant efficiency, p/s) in organs using Living Image software, reflecting LNP biodistribution. Data are represented as mean ± SD (n=5).

Representative bioluminescence imaging (Fig. 3b) revealed variations in fLuc mRNA expression across key organs and tumors, quantified as total luminescence flux in photons per second (p/s). Among the formulations tested, LNP13 exhibited higher expression in the pancreas, while LNP46 showed enhanced expression in the tumor. LNP4 and LNP5 demonstrated notable expressions in the liver. Additionally, LNP13 and LNP46 were observed to express mRNA in the spleen, with LNP13 also showing expression in the lung (Fig. 3c). These findings indicate diverse targeting and efficacy among the LNP formulations, with distinct preferences for tumor, pancreas, and liver accumulation.

RhB fluorescence imaging provided insights into the distribution patterns of LNPs within different organs. Notably, LNP32 showed pronounced accumulation in the tumor, while LNP46 demonstrated distribution towards the liver (Fig. 3d). Quantitative analysis of the fluorescence intensity (total radiant efficiency, p/s) confirmed these observations, highlighting the preferential distribution of certain formulations to targeted organs (Fig. 3e). These findings indicate that specific LNP formulations can be optimized for targeted mRNA delivery, achieving varied expression and biodistribution profiles.

Further quantification of fLuc mRNA bioluminescence and RhB fluorescence was showed for all 48 LNP formulations, grouped into categories based on intensity levels: low (Fig. 4a,4d), medium (Fig. 4b, 4e), and high (Fig. 4c, 4f). This grouping revealed significant formulation-dependent differences in biodistribution (Fig. 4d-4f) and mRNA expression (Fig. 4a-4c) patterns across tumor and major organs, such as the pancreas, spleen, liver, lung, kidney, heart, and colon. In this study, specific LNP formulations such as LNP2, 10, 21, 24, 27, 47 (Fig. 4a) LNP9, 22, 23, 25, 28, 38, 44 (Fig. 4b) and LNP14, 40, 41, 46, 48 (Fig. 4c) demonstrated noteworthy mRNA expression particularly in targeting tumor tissues. These formulations exhibited significant mRNA expression, suggesting their effectiveness in delivering therapeutic mRNA accurately and efficiently. On the other hand, LNP2, 18, 19, 20 (Fig. 4a) L 17, 28 (Fig. 4b) and LNP4, 5, 6, 26, 40 (Fig. 4c) showed pronounced mRNA expression in liver, possibly due to its tailored lipid composition that enhances cellular uptake in liver. Additionally, formulations such as LNP 19, 20, 21, 27, 36 (Fig. 4a), LNP 8, 9, 15, 22, 23, 25, 35, 37,38, 39, 44 (Fig. 4b) and LNP13, 14, 26, 32, 40, 41, 42, 43 46, 48 demonstrated effective mRNA expression in pancreas. LNP10, 21, 24 (Fig. 4a), LNP15, 23, 35 (Fig. 4b) and LNP14 (Fig. 4c) showed some extent of mRNA expression in lung. Furthermore, LNP19, 27, 47 (Fig. 4a), LNP7, 8, 15, 22, 33 (Fig. 4b) and LNP13, 14, 26, 40 (Fig. 4c) showed some extent of mRNA expression in spleen. This clear distinction in performance among the LNPs underscores the critical role of specific formulation characteristics in achieving desired biodistribution and mRNA expression in targeted organs. In the analysis of LNP biodistribution, the graphs (Fig. 4d-f) did not reveal significant differences among the formulations.

**Fig. 4.**
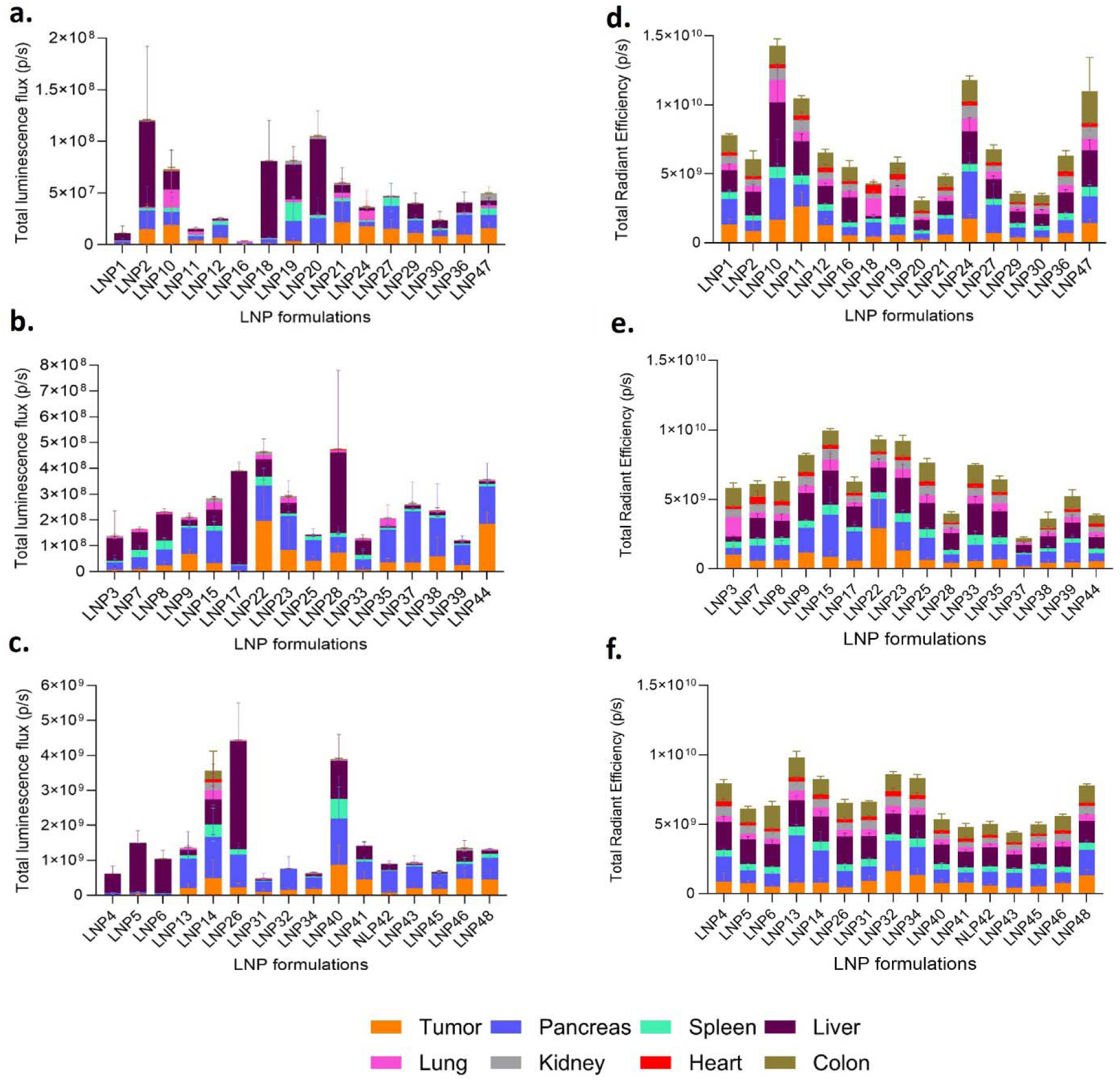
Quantification of Fluc mRNA bioluminescence and RhB fluorescence in individual organs for all 48 LNP formulations grouped into high, medium, and low intensity. (a-c) Total bioluminescence flux (p/s) representing Fluc mRNA expression in major organs (tumor, pancreas, spleen, liver, lung, kidney, heart, and colon) for LNP formulations grouped based on mRNA bioluminescence intensity levels: (a) low, (b) medium, and (c) high. Data was quantified using Living Image software. (d-f) Total radiant efficiency (p/s) representing RhB fluorescence intensity (LNP biodistribution) in major organs for LNP formulations grouped same LNP formulations grouped according to their Fluc bioluminescence levels. Data was quantified using Living Image software. Error bars indicate mean ± SD (n=5). The analysis highlights formulation-dependent differences in biodistribution and mRNA expression across tumor and major organs.

### Lipid compositions and their impact on LNP size

A comprehensive statistical analysis was performed to examine the correlations among various LNP formulation composition and their impact on biophysical characteristics and biological outcomes. The analysis was visualized in a schematic representation (Fig. 5).

**Fig. 5.**
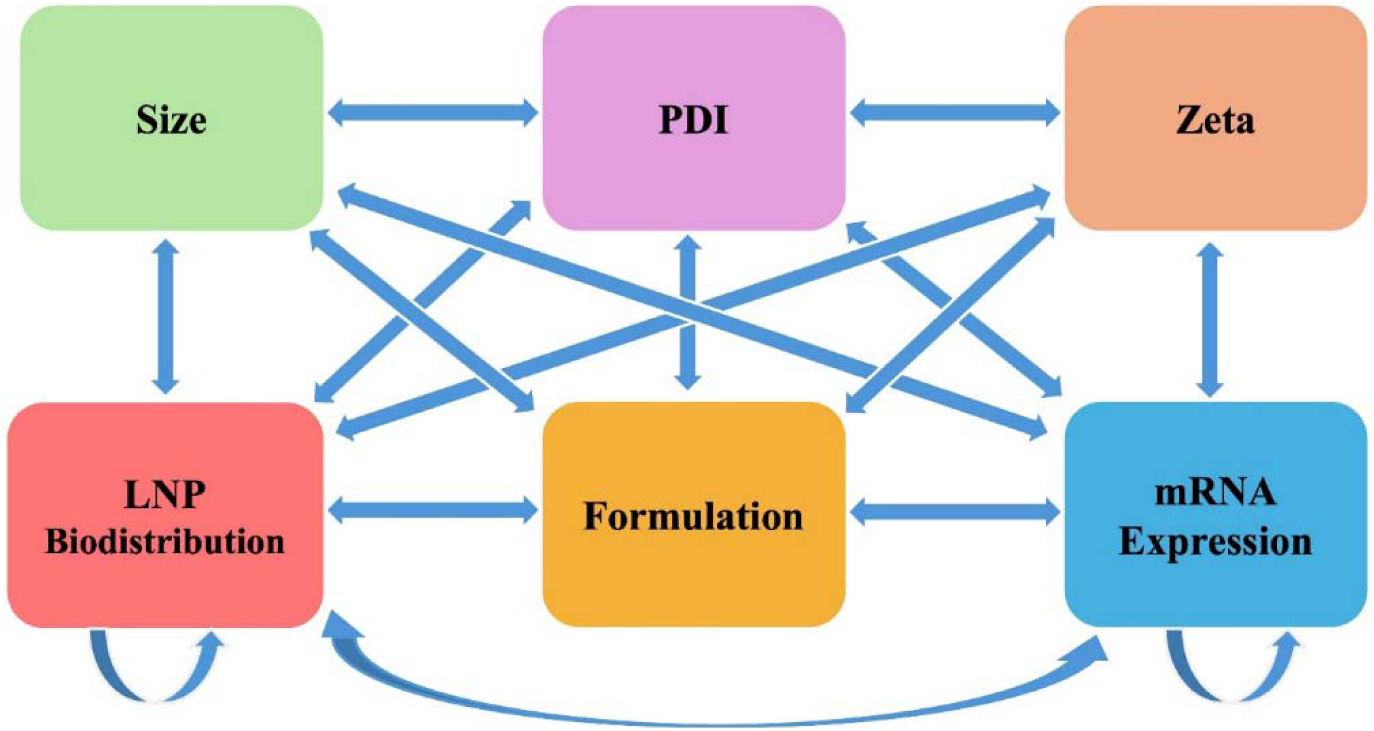
Schematic representation of statistical analysis among formulation parameters and outcomes. The figure illustrates the interrelationships among the lipid compositions and outcomes such as **s**ize, PDI (polydispersity index), zeta potential, LNP biodistribution, and mRNA Expression, as analyzed using JMP software. The schematic highlights the role of **formulation** composition as a central variable influencing physicochemical properties including LNP size, zeta potential, PDI, and downstream outcomes such as biodistribution and mRNA expression. Additionally, correlations between biodistribution and mRNA expression emphasize their interdependency in assessing LNP performance.

Our study employed a robust statistical and predictive analysis framework to elucidate the effects of various lipid compositions on the size of LNPs. Through a detailed exploration involving Kruskal-Wallis tests, ANOVA, decision tree analysis, and predictive modeling, we systematically assessed how different lipid type influences LNP size.

The statistical analysis, using Kruskal-Wallis test and ANOVA, revealed significant effects of ionizable lipids and phospholipids on LNP size, with p-values <0.0001, indicating a strong influence of these lipid types on nanoparticle dimensions. Specifically, FTT5 and DSPC were associated with larger LNP sizes, while GO-C14 and DOPE were associated with smaller LNP sizes. These findings suggest that lipid selection plays a critical role in controlling LNP size (Fig. 6a).

**Fig. 6.**
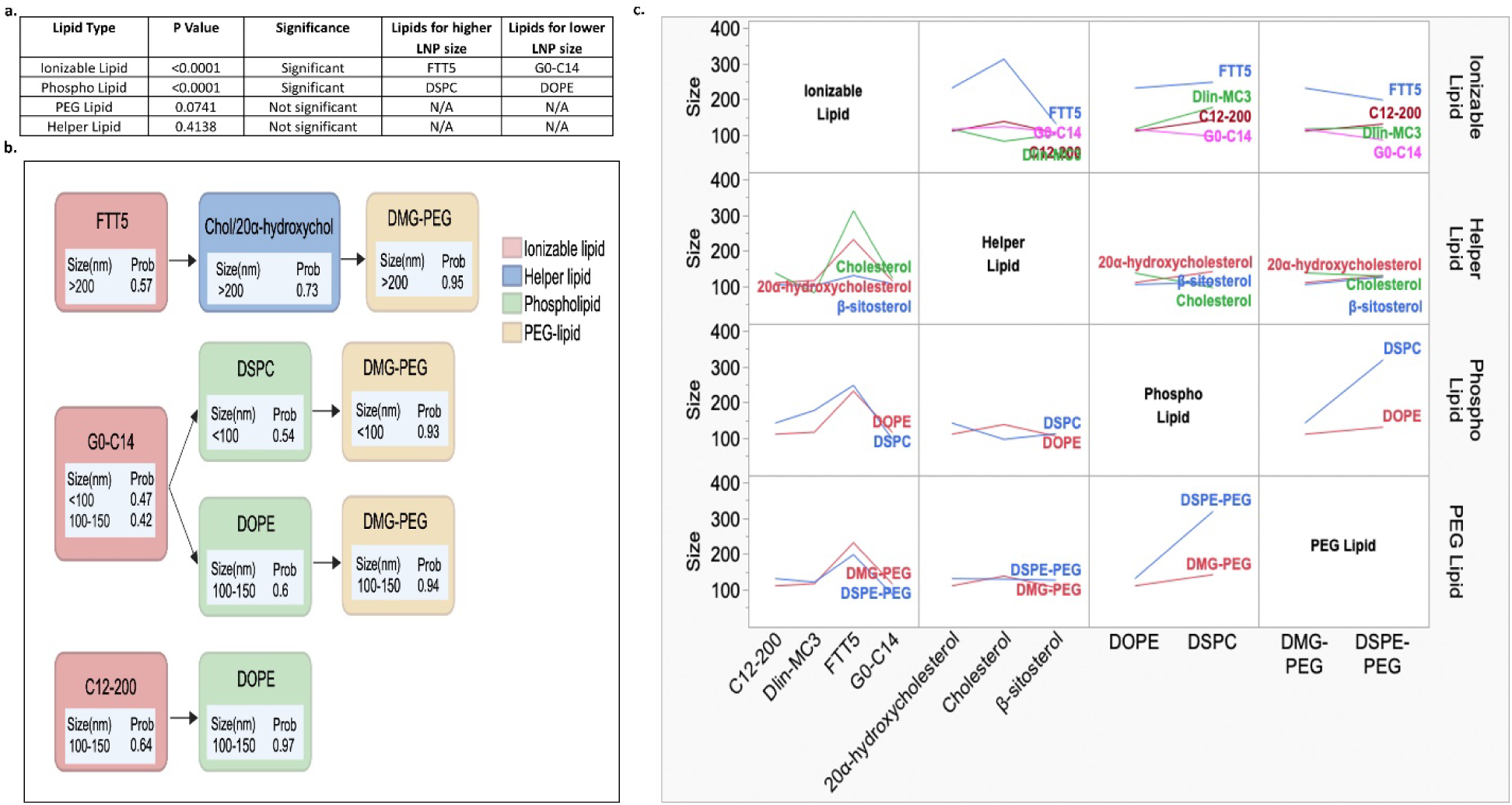
Statistical analysis, decision tree, and predictive modeling analysis for lipid compositions influencing LNP size. (a) Summary table of statistical analysis results, including Kruskal-Wallis test and ANOVA, evaluating the effect of different lipid types on LNP size. The table includes P-values, statistical significance, and the lipids associated with higher and lower LNP sizes for each lipid type. (b) The decision tree, generated using JMP’s Partition method, identifies key lipid compositions influencing LNP size. LNP sizes are categorized as <100 nm, 100–150 nm, 150–200 nm, and >200 nm. Nodes represent lipid components with predicted size categories and probabilities. Lipids are color-coded: ionizable (red), helper (blue), phospholipids (green), and PEG-lipids (yellow). (c) SVR models were trained to capture the relationship between lipid composition and LNP size. Interaction profiles generated using JMP software for the selected SVR model to illustrate how different lipid types and their Interactions influence LNP sizes.

Further elucidation through decision tree analysis allowed for effective categorization of LNP sizes into four groups: <100 nm, 100–150 nm, 150–200 nm, and >200 nm. This part of the study identified significant lipid composition patterns correlating with these size categories. It highlighted ionizable lipids as the primary determinants of size distribution, with helper lipids, phospholipids, and PEG lipids also contributing to size variability but to a lesser extent. Compositions featuring the cationic ionizable lipid FTT5, combined with either cholesterol or 20α-hydroxycholesterol and DMG-PEG, were highly likely to produce LNPs larger than 200 nm. Conversely, LNPs formulated with G0-C14, DSPC, and DMG-PEG typically resulted in sizes under 100 nm. Combinations of G0-C14 with DOPE and DMG-PEG, as well as C12-200 with DOPE yielded LNPs within the 100-150 nm range (Fig. 6b).

The results of the SVR model used to predict the size of LNPs based on lipid formulation features. A grid search optimization experiment refined the model, achieving a high accuracy in predicting LNP size with a Training *R²* of 0.97 and a Validation *R²* of 0.98, demonstrating the robustness of the model. Interaction profiles visualized in JMP highlighted the influence of lipid types and their interactions on LNP size, providing valuable insights into the formulation’s role in LNP size determination. From the interaction profile image (Fig. 6c), FTT5 consistently results in larger LNP sizes when interacting with other lipids, while G0-C14 is associated with smaller LNP sizes across interactions. Among the helper lipids, cholesterol demonstrates the highest LNP sizes, particularly in combination with FTT5. For phospholipids and PEG lipids, the combination of DSPC and DSPE-PEG produces the largest LNP sizes, whereas the pairing of DOPE and DMG-PEG results in smaller LNP sizes. These observations highlight the critical role of lipid combinations in influencing nanoparticle size and provide valuable guidance for optimizing lipid formulations.

### Influence of lipid compositions on LNP zeta potential

Our study conducted a detailed analysis to investigate the effects of various lipid compositions on the zeta potential of LNPs. This investigation combined statistical analysis, decision tree analysis, and predictive modeling to examine how different types of lipids influence the surface charge characteristics of LNPs.

The statistical evaluation included Kruskal-Wallis tests and ANOVA, which identified significant variations in zeta potential attributed to different lipid categories. Ionizable lipids was particularly influential (p-value <0.0001), showing statistically significant differences in zeta potential values. Specific lipid types such as FTT5 was associated with higher zeta potentials, while G0-C14 led to lower zeta potentials. These results are summarized in Fig. 7a, illustrating the role of lipid type in dictating zeta potential characteristics.

**Fig. 7.**
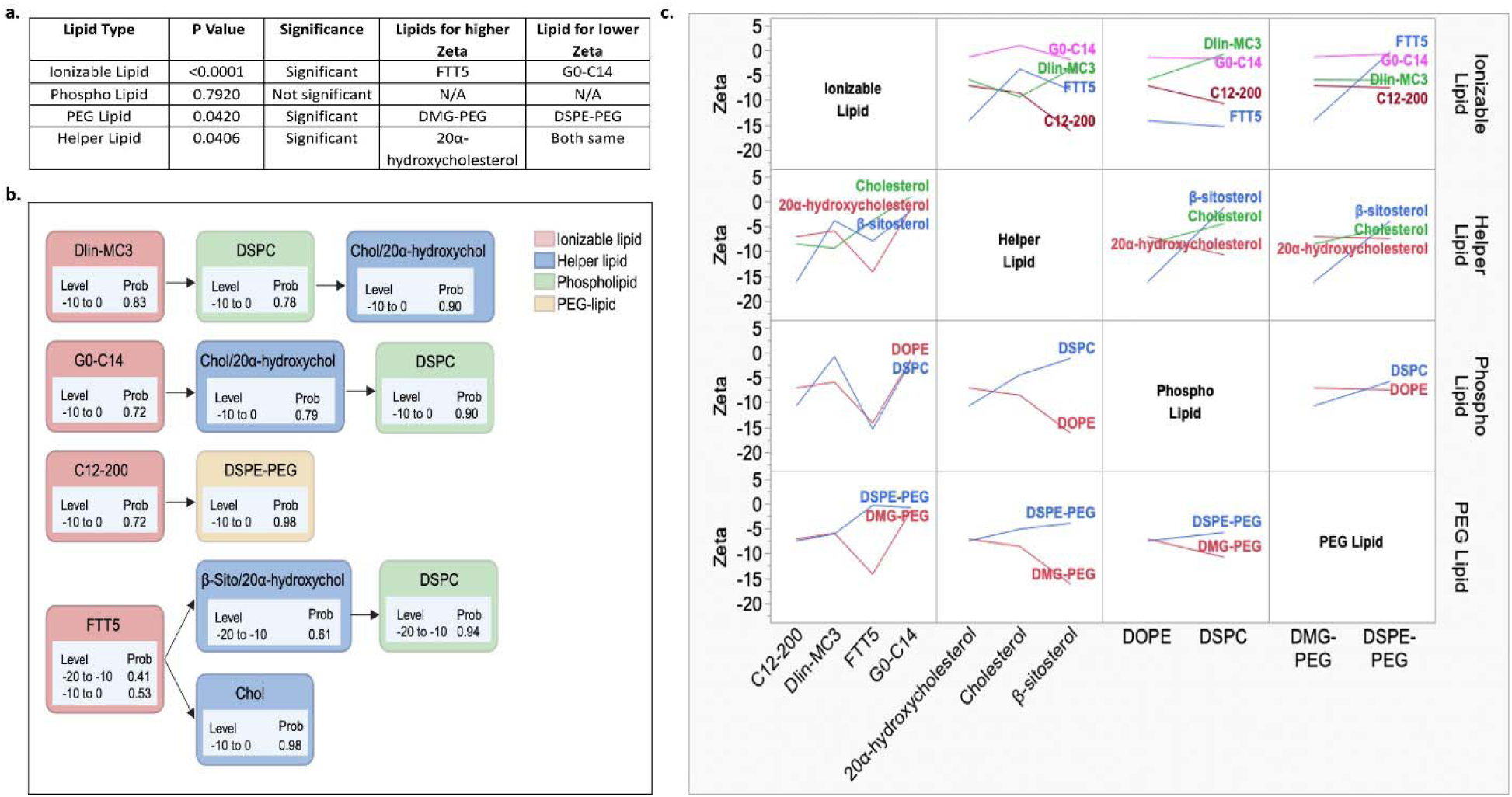
Statistical analysis, decision tree, and predictive modeling analysis for lipid compositions influencing LNP zeta potential. (a) Summary table of statistical analysis results, including Kruskal-Wallis test and ANOVA, evaluating the effect of different lipid types on LNP zeta potential. The table includes P-values, statistical significance, and the lipids associated with higher and lower LNP zeta potential for each lipid type. (b) The decision tree, generated using JMP’s Partition method, identifies key lipid compositions influencing LNP zeta potential. LNP zeta potential is categorized as −20 to −10, −10 to 0, and 0 to 5. Nodes represent lipid components with predicted zeta potential categories and probabilities. Lipids are color-coded: ionizable (red), helper (blue), phospholipids (green), and PEG-lipids (yellow). (c) SVR models were trained to capture the relationship between lipid composition and LNP zeta potential. Interaction profiles generated using JMP software for the selected SVR model to illustrate how different lipid types and their Interactions influence LNP zeta potential.

In the decision tree analysis, the zeta potential was categorized into three groups: −20 to −10, −10 to 0, and 0 to 5 mV. The decision tree effectively mapped the relationship between lipid compositions and zeta potential categories. Notably, lipid combinations such as DLin-MC3-DMA with DSPC and either cholesterol or 20α-hydroxycholesterol, G0-C14 with cholesterol or 20α-hydroxycholesterol and DSPC, C12-200 with DSPE-PEG, and FTT5 with cholesterol all exhibited zeta potentials ranging from −10 to 0 mV. Conversely, formulations like FTT5 with β-sitosterol or 20α-hydroxycholesterol and DSPC were associated with more negative zeta potentials, from −20 to −10 mV. These findings are depicted in Fig. 7b, which illustrates the impact of specific lipid types on the zeta potential values.

Predictive modeling was employed using a SVR with an RBF kernel. The optimization process involved a grid search to refine the model parameters, aiming to minimize prediction errors and maximize the validation *R².* This model demonstrated high accuracy (Training *R²* of 0.91 and a Validation *R²* of 0.92) in predicting zeta potential based on lipid compositions. Interaction profiles visualized in JMP highlighted the influence of lipid types and their impact on zeta (Fig. 7c). From the interaction profile image, FTT5 consistently resulted in more negative zeta potential when interacting with other lipids, while G0-C14 is associated with lower zeta potential across interactions. Helper lipids with DSPE-PEG demonstrated lower zeta potential than DMG-PEG. β-sitosterol demonstrated the higher zeta potential with DOPE than DSPC and with DMG-PEG than DSPE-PEG. These observations highlight the critical role of lipid composition in influencing zeta potential of LNP formulations and elucidate the specific contributions of ionizable, helper, and phospholipids to the surface charge characteristics of LNPs. This comprehensive analysis provides critical insights into how lipid compositions affect the zeta potential of LNPs, influencing their stability and interaction with biological environments.

### Influence of lipid compositions on LNP PDI

Our study employed a robust statistical and predictive analysis framework to elucidate the effects of various lipid compositions on the homogeneity of LNPs. Through a detailed exploration involving Kruskal-Wallis tests, ANOVA, decision tree analysis, and predictive modeling, we systematically assessed how different lipid types of influence homogeneity of LNP formulations.

The statistical analysis, using Kruskal-Wallis test and ANOVA, highlighted PEG-lipid (p-value <0.0001) as the primary determinants of PDI, with ionizable lipids and phospholipids, also contributing to PDI variability but to a lesser extent. Specifically, C12-200, DSPC and DMG-PEG were associated with higher PDI, while FTT5, G0-C14, DOPE and DSPE-PEG were associated with lower PDI. These findings suggest that lipid selection plays a critical role in controlling PDI (Fig. S2a).

Further elucidation through decision tree analysis allowed effective categorization of PDI into three groups: <0.1, 0.1-0.2, and >0.2. This part of the study identified significant lipid composition patterns correlating with these PDI categories. Combinations of G0-C14 with DMG-PEG and DOPE were linked to PDIs 0.1- 0.2. Formulations containing FTT5 or DLin-MC3-DMA with DSPE-PEG, paired with either β-sitosterol or 20α-hydroxycholesterol, consistently resulted in PDIs < 0.1. Conversely, C12-200 combined with DSPC and either β-sitosterol or 20α-hydroxycholesterol typically produced PDIs greater than 0.2. (Fig. S2b).

Predictive modeling using a SVR with an RBF kernel was then applied. A grid search optimization experiment refined the model, achieving a high accuracy in predicting PDI with a Training *R²* of 0.88 and a Validation *R²* of 0.89, demonstrating the robustness of the model. Interaction profiles visualized in JMP highlighted the influence of lipid types and their role in PDI determination. Interaction profiles generated using JMP software visually represented the complex interplay between different lipid types and their concentrations, impacting the homogeneity. From the interaction profile image (Fig. S2c), DLin-MC3-DMA with any helper lipids with DOPE and DMG-PEG consistently resulted in higher PDI. In contrast, FTT5 with any of the three helper lipids or DOPE resulted in lower PDI and DSPC higher PDI.

### LNP biodistribution influenced by lipid compositions

Our study conducted comprehensive statistical analyses and decision tree evaluations to understand the influence of lipid formulations on the biodistribution of LNPs across various organs. These analyses included ANOVA, Kruskal-Wallis tests, and decision tree methodologies, focusing on the relationship between specific lipid components and the distribution patterns of LNPs.

The ANOVA results showed significant variability in LNP biodistribution across 48 different LNP formulations, particularly in key organs such as the tumor, pancreas, spleen, and liver. This variation is depicted in Fig. 8a and Fig. S3a, which illustrates each data point representing RhB fluorescence intensity from biodistribution studies involving 5 mice per formulation. The results indicate no significant intra-formulation variability but notable differences between formulations. These findings are substantiated by statistically significant differences observed across multiple organs, highlighting the role of formulation in achieving targeted biodistribution. The p-values reported (>0.05) for different major organs and tumor in the five mice for the same formulation showed no significant changes (Fig. 8b, top table), whereas significant inter-formulation variability was confirmed by p-values less than 0.0001 across all major organs and the tumor (Fig. 8b, bottom table).

**Fig. 8.**
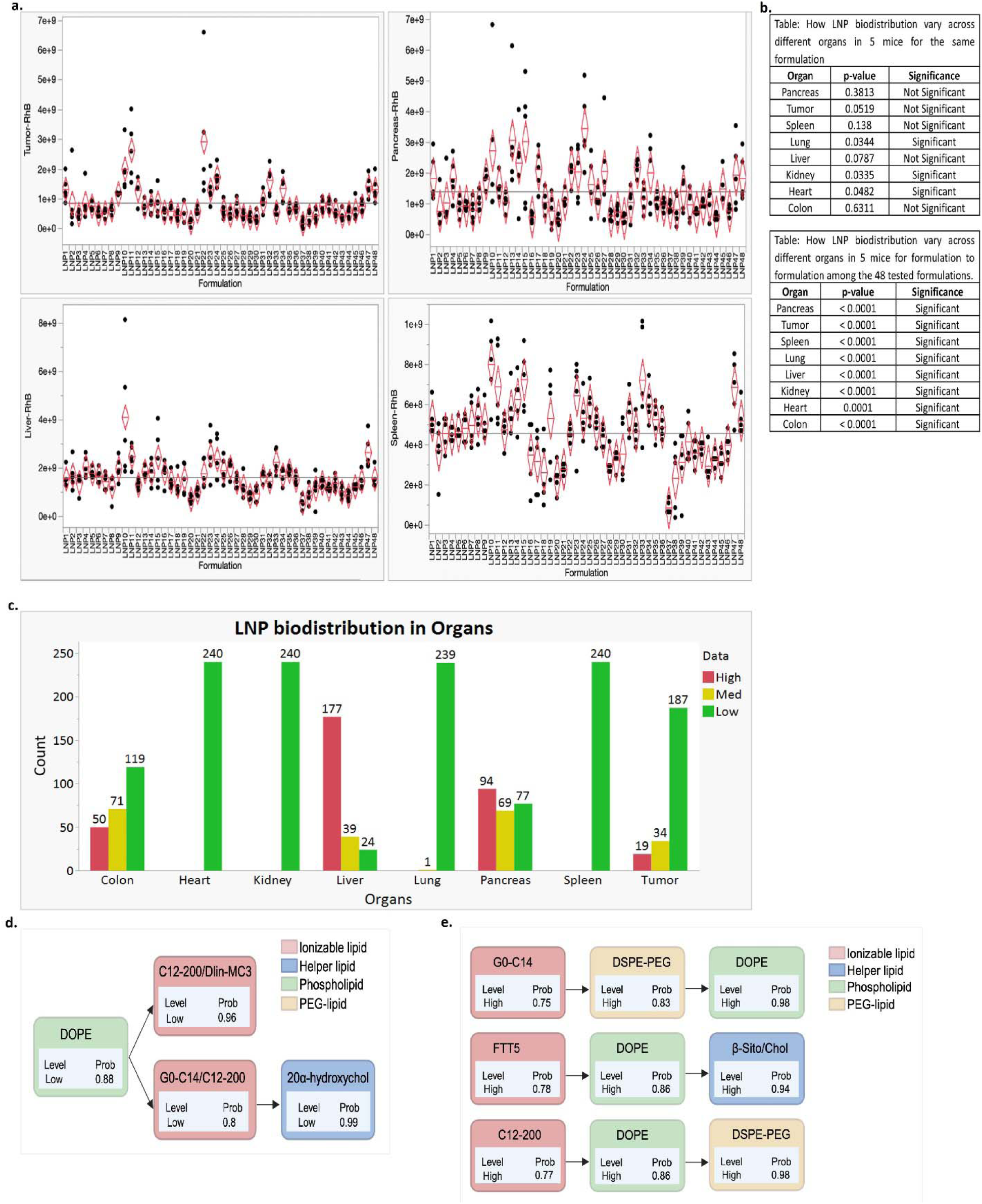
Statistical analysis of LNP biodistribution influenced by formulation composition across different organs. (a) ANOVA results showing LNP biodistribution (RhB fluorescence intensity) across 48 formulations in tumor (top left), pancreas (top right), liver (bottom left) and spleen (bottom right). The y-axis represents biodistribution levels, while the x-axis represents the LNP formulations. Each column represents data from five mice for the same formulation, showing minimal variability within the same formulation but significant differences across different formulations. (b) Results of the ANOVA test examining the variability of biodistribution. The top table shows the variability across the same formulation for different mice. The bottom table highlights variability across different formulations. (c) Categorization of LNP biodistribution levels into high (red), medium (yellow), and low (green) across organs. The bar graph shows the count of formulations achieving high, medium, or low biodistribution level for each organ. (d, e) Decision tree analysis categorized normalized RhB fluorescence intensity into high, medium, and low levels based on formulation compositions. Using the partition method in JMP software, the trees identified relationships between lipid components and RhB fluorescence intensity. (d) shows the analysis for tumor biodistribution, and (e) shows the analysis for liver biodistribution. The trees were pruned to retain the most significant split rules, focusing on lipid components with strong predictive power for high or low RhB fluorescence intensity. Nodes represent lipid components with predicted intensity categories and probabilities. Lipids are color-coded: ionizable (red), helper (blue), phospholipids (green), and PEG-lipids (yellow).

We categorized the biodistribution levels into high, medium, and low categories for each organ studied. As shown in Fig. 8c, organs like the liver, pancreas, colon and tumor frequently exhibited higher (red) biodistribution levels across multiple formulations. In contrast, other organs like the heart, lung, kidney and spleen showed lower (green) distribution levels, which helps in identifying which organs are more likely to accumulate LNPs.

The Kruskal-Wallis test (Table S2) revealed significant differences in RhB fluorescence intensity across various organ tissues, indicating significant variability in the biodistribution of LNPs, which can be attributed to differences in lipid composition. Statistical analysis using the Kruskal-Wallis test and ANOVA confirmed significant differences in the distribution patterns among different types of lipids within each lipid category. In the pancreas, significant differences in RhB biodistribution intensity were observed among ionizable lipids (p-value: 0.0054) and PEG lipids (p-value: 0.0099) with significant distribution for DLin-MC3-DMA, G0-C14 and DMG-PEG respectively. In the tumor, significant differences were noted for phospholipids (p-value: <0.0001) and PEG lipids (p-value: <0.0001) with significant distribution for DSPC and DSPE-PEG respectively. In the liver, significant differences were found for ionizable lipids (p-value: 0.0211) with FTT5 being the most effective, and in the spleen, significant differences were also observed for ionizable lipids (p-value: 0.0035) with significant distribution for FTT5. These results highlight the critical role of lipid composition in influencing LNP biodistribution, suggesting the potential for optimizing LNP formulations for targeted organ delivery.

The decision tree analysis offered detailed insights into the impact of specific lipid compositions on LNP biodistribution in the tumor (Fig. 8d), liver (Fig. 8e), and pancreas (Fig. S3b). This approach classified normalized RhB fluorescence intensities into three levels high, medium, and low based on the lipid components. To focus on the most influential factors, the trees were pruned to highlight lipid components that strongly predict different biodistribution levels. The analysis indicated that combinations such as DOPE with DLin-MC3-DMA or FTT5, and DOPE with G0-C14 or C12-200 with 20α-hydroxycholesterol, were associated with low LNP distribution to the tumor. Conversely, formulations like G0-C14 with DSPE-PEG and DOPE, FTT5 with DOPE and either cholesterol or β-sitosterol, and C12-200 with DOPE and DSPE-PEG showed high LNP distribution to the liver. Formulations of G0-C14 with 20α-hydroxycholesterol and DMG-PEG, G0-C14 with cholesterol and DMG-PEG, and DLin-MC3-DMA with DSPC and 20α-hydroxycholesterol were associated with high LNP biodistribution in the pancreas. In contrast, a combination of C12-200 with DOPE and β-sitosterol resulted in low LNP biodistribution in the pancreas.

### Influence of lipid compositions on mRNA expression levels in organs

This section of the study examines the impact of different LNP formulations on mRNA expression across various organs, utilizing comprehensive statistical methods and decision tree analyses. These analyses help elucidate how specific lipid compositions can optimize mRNA delivery to targeted tissues.

The statistical examination ANOVA test was applied to assess mRNA expression across 48 different LNP formulations in major organs such as the tumor, pancreas, spleen, and liver. This variation is depicted in Fig. 9a and Fig. S4a, which illustrates each data point representing mRNA expression intensity from studies involving 5 mice per formulation. The results indicated that mRNA expression varied significantly across different formulations but showed no significant variability within the same formulation when tested across five mice. This points to the critical influence of LNP formulation on mRNA expression efficiency. These findings are substantiated by statistically significant differences observed across multiple organs, highlighting the role of formulation in achieving targeted mRNA expression. The p-values reported (>0.05) for different major organs and tumor in the five mice for the same formulation showed no significant changes (Fig. 9b, top table), whereas significant inter-formulation variability in pancreas, tumor, spleen, liver and was confirmed by p-values < 0.0001 and lung with p-value 0.0019 (Fig. 9b, bottom table).

**Fig. 9.**
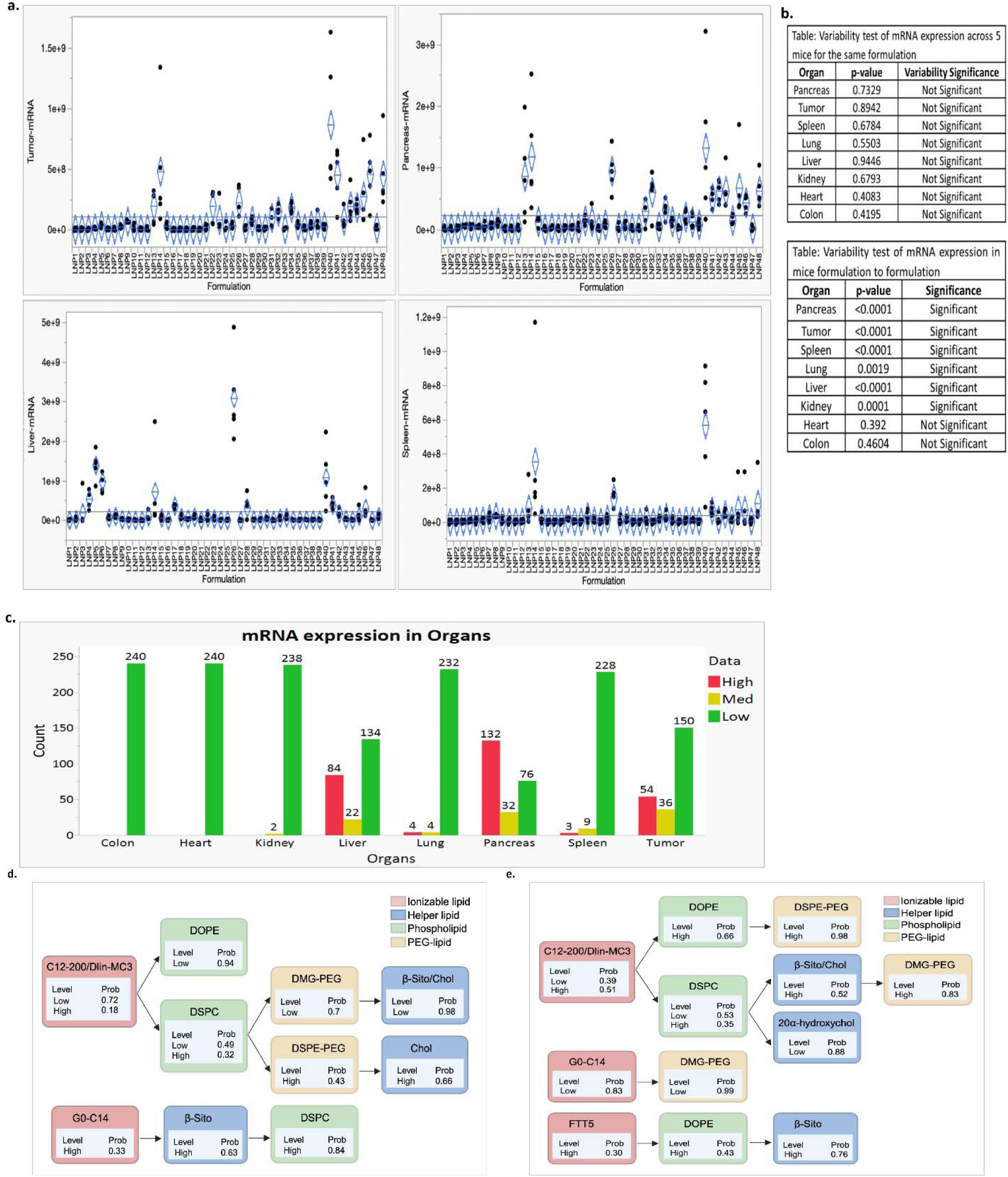
Statistical analysis of mRNA expression levels influenced by LNP formulation composition across different organs. (a) ANOVA results showing mRNA expression across 48 LNP formulations in tumor (top left), pancreas (top right), liver (bottom left) and spleen (bottom right). The y-axis represents mRNA expression levels, while the x-axis represents the LNP formulations. Each column represents data from five mice, demonstrating minimal variability within the same formulation but significant differences across different formulations. (b) Results of the ANOVA test examining the variability of mRNA expression. The top table shows the variability across the same formulation for different mice. The bottom table highlights variability across different formulations. (c) Categorization of mRNA expression levels into high (red), medium (yellow), and low (green) across organs. The bar graph shows the count of formulations achieving three expression levels for each organ. (d, e) Decision tree analysis categorized normalized mRNA expression levels into high, medium, and low based on formulation compositions. Using the partition method in JMP software, the trees identified relationships between lipid components and mRNA expression levels. (d) shows the analysis for tumor mRNA expression, and (e) shows the analysis for liver mRNA expression. The trees were pruned to retain the most significant split rules, focusing on lipid components with strong predictive power for high or low expression levels. Nodes represent lipid components with predicted expression categories and probabilities. Lipids are color-coded: ionizable (red), helper (blue), phospholipids (green), and PEG-lipids (yellow).

Further analysis categorized the mRNA expression levels into high (red), medium (yellow), and low (green) categories across different organs. This categorization, shown in Fig. 9c, highlights the variations in expression efficacy, with certain organs like the tumor, pancreas and liver frequently achieving higher expression levels. While lung, spleen, heart, colon and kidney demonstrated low mRNA expression intensity. This categorization helps identify which organ systems are more effectively targeted by specific LNP formulations, facilitating the design of more efficient gene therapy delivery systems.

Analysis of fLuc bioluminescence across various organs highlighted the influence of lipid type on the efficiency and localization of LNP-mediated gene delivery. Significant variations in bioluminescence intensities across different lipid types were documented, showing varying degrees of efficacy depending on the organ, as confirmed by Kruskal-Wallis and ANOVA tests (Table S3). In the pancreas, ionizable lipids had the most pronounced effect on fLuc bioluminescence intensity, with GO-C14 demonstrating the highest efficiency in mRNA expression (p-value <0.0001), whereas C12-200 was identified as the least effective. Notable impacts were also seen with phospholipids (p-value: 0.0074) and PEG lipids (p-value: 0.0125), where DSPC and DMG-PEG emerged as the most impactful, respectively, and cholesterol was identified as the most effective helper lipid (p-value: 0.0710), while 20α-hydroxycholesterol was the least. In tumor tissues, significant results were recorded for ionizable and phospholipid types (p < 0.0001), with GO-C14 and DSPC being the top performers. However, helper lipids showed no significant effect on bioluminescence in tumors (p-value: 0.9305). In the spleen, DSPC as a phospholipid significantly enhanced bioluminescence expression (p-value: 0.0032). In the liver, substantial differences were noted across all lipid types except helper lipids, with ionizable lipid C12-200, phospholipid DOPE, and PEG lipid DSPE-PEG showing the most significant enhancement in bioluminescence visibility and intensity (p-values < 0.0001 and 0.0004, respectively).

Decision tree analysis was conducted to further detail the relationship between lipid composition and mRNA expression levels in the tumor (Fig. 9d), liver (Fig. 9e) and pancreas (Fig. S4b). This analysis depicts how the mRNA expression levels were grouped into high, medium, and low based on specific lipid compositions. Using JMP-based decision tree analysis, the study identified specific lipid compositions that significantly dictate organ-specific mRNA expression. High tumor expression was primarily associated with formulations containing the ionizable lipid G0-C14 combined with β-sitosterol and DSPC. In contrast, tumor expression remained low when using C12-200 or DLin-MC3-DMA in combination with DOPE, DMG-PEG, and either cholesterol or β-sitosterol. Liver-specific targeting followed different rules: high mRNA expression was linked to C12-200 or DLin-MC3-DMA when formulated with DOPE and DSPE-PEG, or when combined with DSPC, DMG-PEG, and either β-sitosterol or cholesterol. High liver expression was also noted for FTT5 with DOPE and β-sitosterol. Conversely, the combination of G0-C14 with DMG-PEG and 20α-hydroxycholesterol resulted in low liver expression. Notably, the G0 C14/DSPC/DSPE PEG LNP emerged as a lead candidate, achieving a > 6 fold increase in tumor luciferase signal relative to the library median while simultaneously reducing liver exposure by ∼60%. In the pancreas, high expression was associated with several specific combinations: G0-C14 with 20α-hydroxycholesterol; G0-C14 with cholesterol and DMG-PEG; FTT5 with DSPC and DSPE-PEG; and DLin-MC3-DMA with DMG-PEG and DOPE. However, C12-200 combined with DOPE and DSPE-PEG resulted in low pancreatic expression. By pruning the decision trees to focus on these statistically significant components, the analysis provides a clear roadmap of how individual lipids drive effective, site-specific mRNA delivery.

### Pairwise correlation analysis of RhB intensities and mRNA expression across various organs

In our study, the correlation analysis between RhB tracking (LNP biodistribution) and mRNA expression levels was instrumental in identifying key interrelationships essential for refining LNP formulations for targeted gene delivery. Notably, a moderate negative correlation (r = −0.4024) between RhB distribution in the pancreas and liver suggested that higher RhB levels in the pancreas are associated with reduced levels in the liver, indicating an inverse relationship. Conversely, the correlations between RhB distributions in the pancreas and tumor (r = 0.1718) and between the liver and tumor (r = −0.1311) were found to be very weak and statistically insignificant, suggesting negligible influence of RhB distribution among these organs (Table S4 top). For mRNA expression patterns, a weak positive correlation (r = 0.2790) between mRNA levels in the pancreas and tumor suggested a slight positive association, where increased mRNA expression in the pancreas may marginally elevate expression in the tumor. In stark contrast, there was a strong negative correlation (r = −0.6896) between mRNA expression in the pancreas and liver, indicating that higher mRNA expression in the pancreas strongly correlates with decreased expression in the liver. Furthermore, mRNA expression between the liver and tumor also showed a moderate negative correlation (r = −0.4204), suggesting that increased mRNA expression in the liver tends to coincide with reduced expression in the tumor (Table S4 middle). Additionally, the correlation analysis within the same organs revealed varied strengths of positive associations between mRNA expression and RhB biodistribution. In the pancreas, a moderate positive correlation (r = 0.3305) suggested that increased mRNA expression is moderately associated with higher RhB biodistribution. In the tumor, mRNA and RhB levels also demonstrated a moderate positive correlation (r = 0.4798), indicating that effective RhB distribution is correlated with enhanced mRNA delivery in tumor tissues. However, in the liver, only a weak positive correlation (r = 0.2534) was observed, indicating a limited association between RhB distribution and mRNA expression (Table S4 bottom). This comprehensive analysis underscores the complex dynamics governing the interactions of LNP components in different organ environments, highlighting the importance of targeted formulation adjustments for improved gene delivery efficacy.

### Influence of LNP size, zeta and PDI on LNP biodistribution and mRNA expression levels in organs

Our pairwise correlation analysis of particle size (Table S5 top), zeta potential (Table S5 bottom), and PDI (Table S7), with mRNA expression levels and LNP biodistribution, unveiled interrelationships critical for optimizing LNP formulations for targeted gene delivery. We found a weak positive correlation between LNP size and biodistribution (Pearson’s r = 0.2467) and between particle size and mRNA expression levels (Pearson’s r = 0.2816), indicating that larger LNPs tend to be more effective in delivering mRNA to tumors. Conversely, a very weak negative correlation between size and mRNA expression in the liver (Pearson’s r = −0.1763) suggests that smaller particles enhance mRNA expression in the liver. The analysis also showed no correlation between zeta potential and LNP biodistribution and a very weak positive correlation between zeta potential and mRNA expression in tumors (Pearson’s r = 0.2103), suggesting that a higher negative charge may improve mRNA expression in tumors. Furthermore, no correlation was observed between formulation uniformity and either LNP biodistribution or mRNA expression. These findings underscore the complex dynamics between the physical attributes of LNPs and their biological efficacy, emphasizing the importance of meticulous design considerations regarding particle size, uniformity, and surface charge in developing effective LNP-based delivery systems.

## DISCUSSION

Our study demonstrates that the formulations employed are non-toxic and emphasizes the significant influence of lipid composition on the properties and efficacy of LNPs in gene delivery applications. Specific lipid combinations significantly affected the size, zeta potential, and mRNA expression profiles of LNPs, key factors that influence their stability, cellular interactions, and functional performance. For instance, formulations containing FTT5 with cholesterol/20α-hydroxycholesterol and DMG-PEG consistently produced larger (>200nm) LNP formulations, provide prolonged circulation time and higher cargo capacity, which are advantageous for therapies requiring sustained release or the delivery of larger genetic payloads. While those with G0-C14, DSPC, and DMG-PEG resulted in smaller (<100nm) nanoparticle sizes, enhance tissue penetration and reduce clearance by the immune system, making them highly effective for systemic delivery and reaching deep tissue targets. In terms of zeta potential, DLin-MC3-DMA / G0-C14 with DSPC and cholesterol/20α-hydroxycholesterol, exhibited smaller zeta potentials (−10 to 0 mV). Conversely, formulations like FTT5 with DSPC and β-sitosterol/20α-hydroxycholesterol were associated with more negative zeta potentials (−20 to −10 mV) enhancing specific cellular interactions or stability. This demonstrates how lipid composition affects surface charge characteristics that are critical for colloidal stability and biological interactions. The mRNA expression profiles varied significantly with different lipid compositions, highlighting the capability of targeted formulations to enhance or suppress gene expression in specific organs. G0-C14 combined with β-sitosterol and DSPC was notably effective in promoting high mRNA expression in tumor tissues, whereas C12-200/ DLin-MC3-DMA with DOPE generally showed lower expression levels. Combinations such as FTT5 with DOPE and β-sitosterol markedly increased mRNA expression in the liver, underscoring the potential for organ-specific gene delivery strategies. Our correlation analysis revealed a negligible or very weak correlation of LNP size and zeta with LNP biodistribution and mRNA expression. A moderate negative correlation between LNP biodistribution in the pancreas and liver and a strong negative correlation between mRNA expression in the pancreas and liver suggests that enhanced LNP biodistribution and mRNA expression in one organ can inversely impact another, which is crucial for the strategic design of LNPs for targeted therapy. Moreover, positive correlations within the pancreas and tumor suggest that effective LNP biodistribution aligns with enhanced mRNA expression, highlighting the potential for co-optimizing these properties to improve therapeutic outcomes.

While this study provides a comprehensive map of the formulation space for IP-administered mRNA-LNPs in pancreatic cancer, several limitations must be acknowledged. First, our analysis of biodistribution and mRNA expression was conducted at a single 12-hour time point. Although this interval is commonly used to capture peak expression for many LNP systems, future studies will involve longitudinal imaging to fully characterize the kinetics of mRNA stability and protein production. Second, these findings are based on the KPC8060 orthotopic tumor model; to ensure broader translational relevance, subsequent work will validate the top-performing formulations in diverse models, including patient-derived xenografts (PDX) that better capture human tumor heterogeneity. Finally, this library screen utilized reporter fLuc-mRNA to optimize delivery efficiency. Future investigations will transition to therapeutic payloads such as mRNA-encoded cytokines or tumor suppressors to evaluate anti-tumor efficacy. Furthermore, the current study utilized a single dose across formulations; dose-response studies are required to define the therapeutic window and optimal dosing frequency. These future efforts will build upon the structure-activity relationships established here to refine LNP design for the effective treatment of pancreatic ductal adenocarcinoma.

## CONCLUSION

In conclusion, our findings affirm the critical role of lipid composition in optimizing the size, zeta potential, and LNP delivery and mRNA expression efficacy of LNPs, offering insights that could significantly advance the development of LNP-based gene therapies. By leveraging these insights, future research can refine LNP formulations to enhance their clinical applicability, particularly for targeted gene expression in organs such as the liver, pancreas, and tumors, thus improving the precision and effectiveness of gene therapies.

## Acknowledgments

The authors would like to thank the Small Animal Imaging Facility at the University of Nebraska Medical Center for their technical support with the IVIS studies. We are also grateful to Ember A. Eldridge from the Tissue Sciences Facility at UNMC for her assistance with animal tissue sectioning and H&E staining, and Dr. Samuel Cohen at UNMC for his analysis of tissue toxicity.

## Author contributions

Conceptualization: D.O., F.I.

Formal analysis: F.I., M.A.

Funding acquisition: D.O.

Investigation: F.I., A.D., M.A., L.D., N.K.

Methodology: F.I.

Project administration: D.O.

Supervision: D.O.

Writing - original draft: F.I.

Writing – review & editing: F.I., M.A., D.O.

## Data and materials availability

All data are available in the main text or the supplementary materials.

