## Supporting Figures and Tables for "Structure-Activity Mapping of Intraperitoneal mRNA-LNPs: Decoupling Tumor and Liver Biodistribution in Pancreatic Cancer"

Farhana Islam *et al.*

**This PDF file includes:**

Figs. S1 to S4
Tables S1 to S7

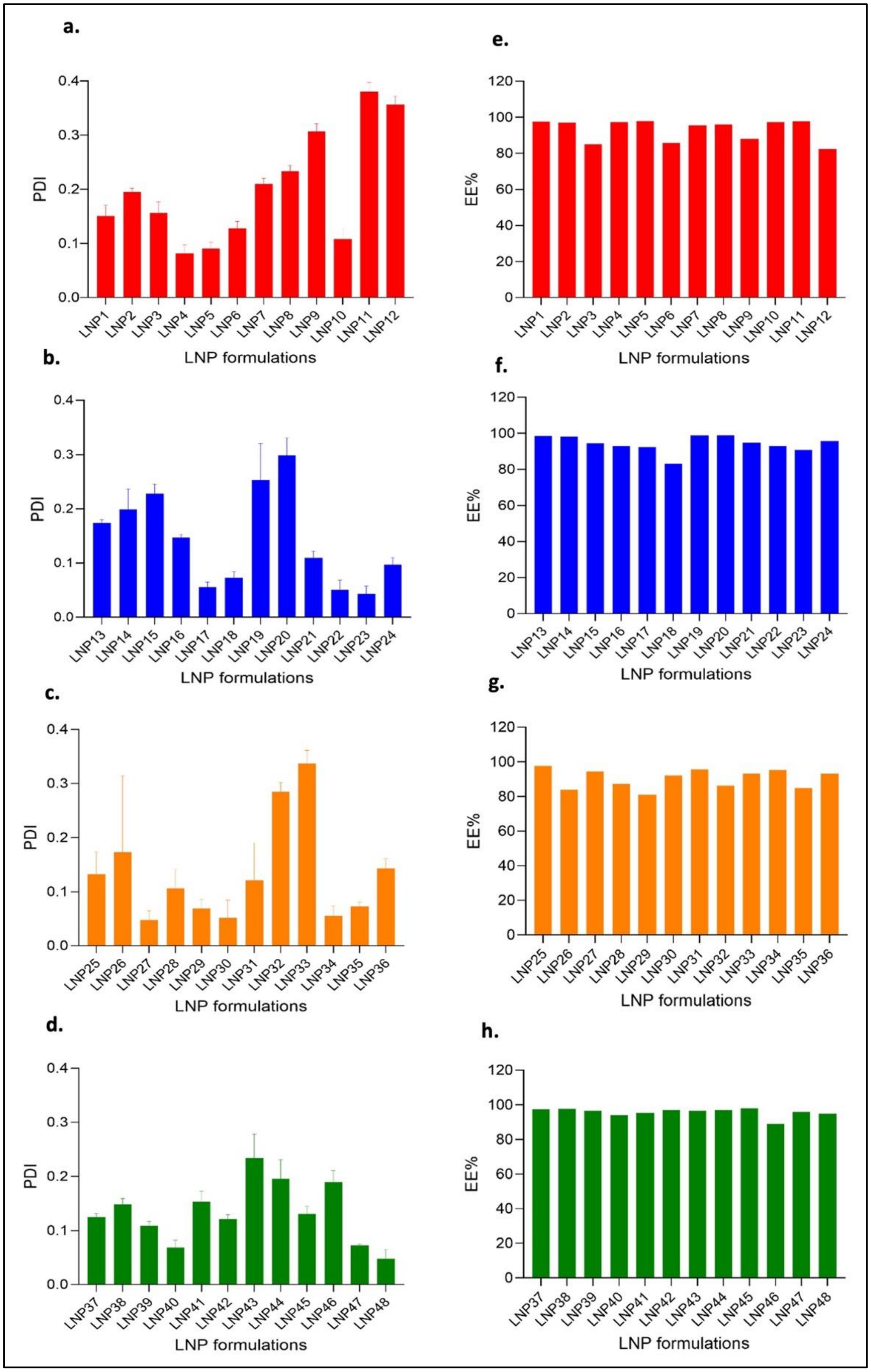

a.
PDI
LNP1 LNP2 LNP3 LNP4 LNP5 LNP6 LNP7 LNP8 LNP9 LNP10 LNP11 LNP12
LNP formulations
b.
PDI
LNP13 LNP14 LNP15 LNP16 LNP17 LNP18 LNP19 LNP20 LNP21 LNP22 LNP23 LNP24
LNP formulations
c.
PDI
LNP25 LNP26 LNP27 LNP28 LNP29 LNP30 LNP31 LNP32 LNP33 LNP34 LNP35 LNP36
LNP formulations
d.
PDI
LNP37 LNP38 LNP39 LNP40 LNP41 LNP42 LNP43 LNP44 LNP45 LNP46 LNP47 LNP48
LNP formulations
e.
EE%
LNP1 LNP2 LNP3 LNP4 LNP5 LNP6 LNP7 LNP8 LNP9 LNP10 LNP11 LNP12
LNP formulations
f.
EE%
LNP13 LNP14 LNP15 LNP16 LNP17 LNP18 LNP19 LNP20 LNP21 LNP22 LNP23 LNP24
LNP formulations
g.
EE%
LNP25 LNP26 LNP27 LNP28 LNP29 LNP30 LNP31 LNP32 LNP33 LNP34 LNP35 LNP36
LNP formulations
h.
EE%
LNP37 LNP38 LNP39 LNP40 LNP41 LNP42 LNP43 LNP44 LNP45 LNP46 LNP47 LNP48
LNP formulations

**Fig. S1. Characterization of 48 mRNA-LNP formulations grouped in four different cationic ionizable lipids.** Each lipid library contains 12 formulations. (a-d) Polydispersity index (PDI) of mRNA-loaded lipid nanoparticles (LNPs) formulated with C12-200 (a, red), DLin-MC3 (b, blue), FTT5 (c, orange), and G0-C14 (d, green), measured via dynamic light scattering (DLS). (e-h) Encapsulation efficiency (EE%) of mRNA-LNP formulations prepared with C12-200 (e, red), DLin-MC3 (f, blue), FTT5 (g, orange), and G0-C14 (h, green), measured via ribogreen assay.

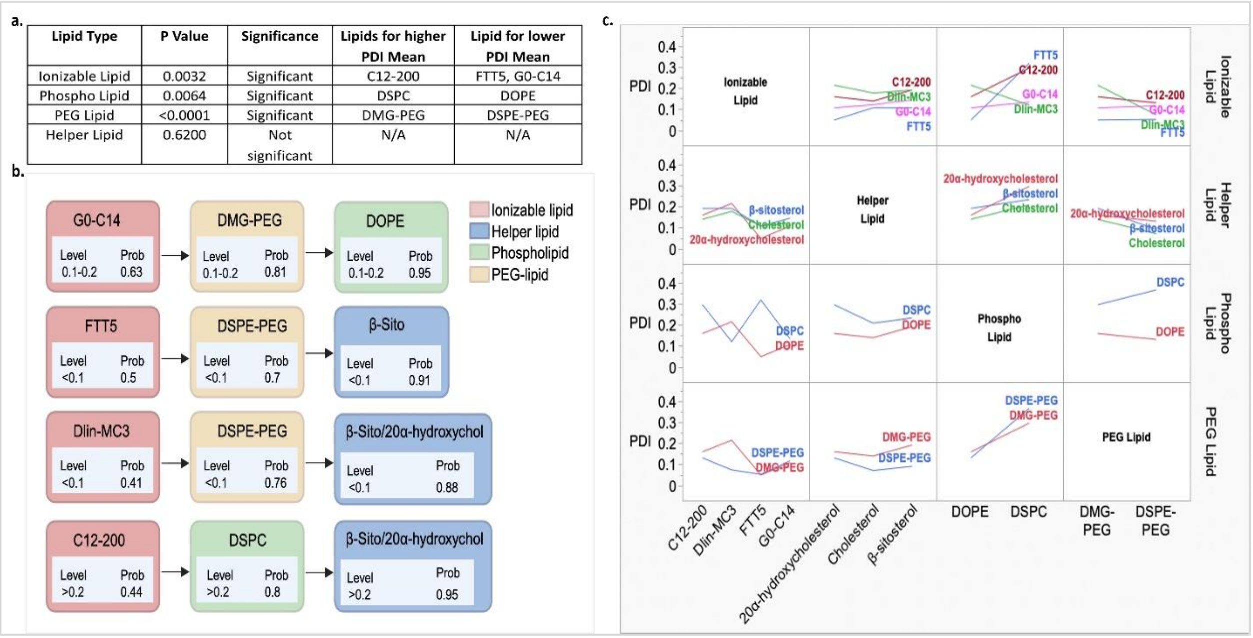


**Fig. S2. Statistical analysis, decision tree, and predictive modeling analysis for lipid compositions influencing LNP homogeneity (PDI).** (a) Summary of Kruskal-Wallis test and ANOVA results analyzing the effect of lipid types (ionizable lipids, phospholipids, PEG lipids, and helper lipids) on LNP homogeneity. (b) The decision tree, generated using JMP's Partition method, identifies key lipid compositions influencing LNP homogeneity. LNP homogeneity (PDI) is categorized as <0.1, 0.1-0.2, and >0.2. Nodes represent lipid components with predicted homogeneity categories and probabilities. Lipids are color-coded: ionizable (red), helper (blue), phospholipids (green), and PEG-lipids (yellow). (c) SVM models were trained to capture the relationship between lipid composition and LNP homogeneity (PDI). Interaction profiles generated using JMP software for the selected SVM model to illustrate how different lipid types and their Interactions influence LNP homogeneity.

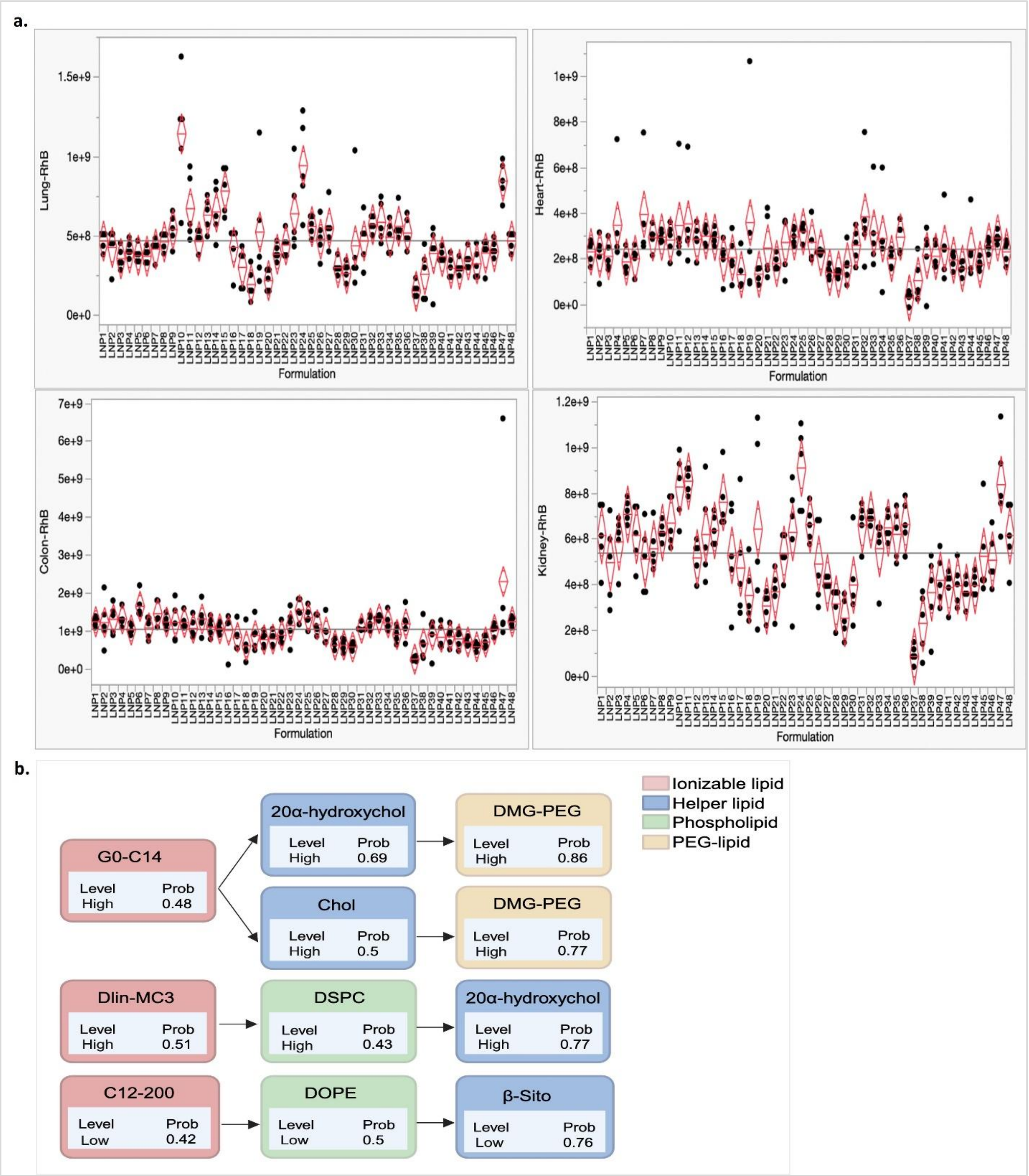


**Fig. S3. Statistical analysis of LNP biodistribution influenced by formulation composition across different organs.** (a) ANOVA results showing LNP biodistribution (RhB fluorescence intensity) across 48 formulations in lung (top left), heart (top right), colon (bottom left) and kidney (bottom right). The y-axis represents biodistribution levels, while the x-axis represents the LNP formulations. Each column represents data from five mice for the same formulation, showing minimal variability within the same formulation but significant differences across different formulations. *(b)* Decision tree analysis categorized normalized RhB fluorescence intensity into high, medium, and low levels based on formulation compositions. Using the partition method in JMP software, the trees identified relationships between lipid components and RhB fluorescence intensity shows the analysis for pancreas biodistribution.

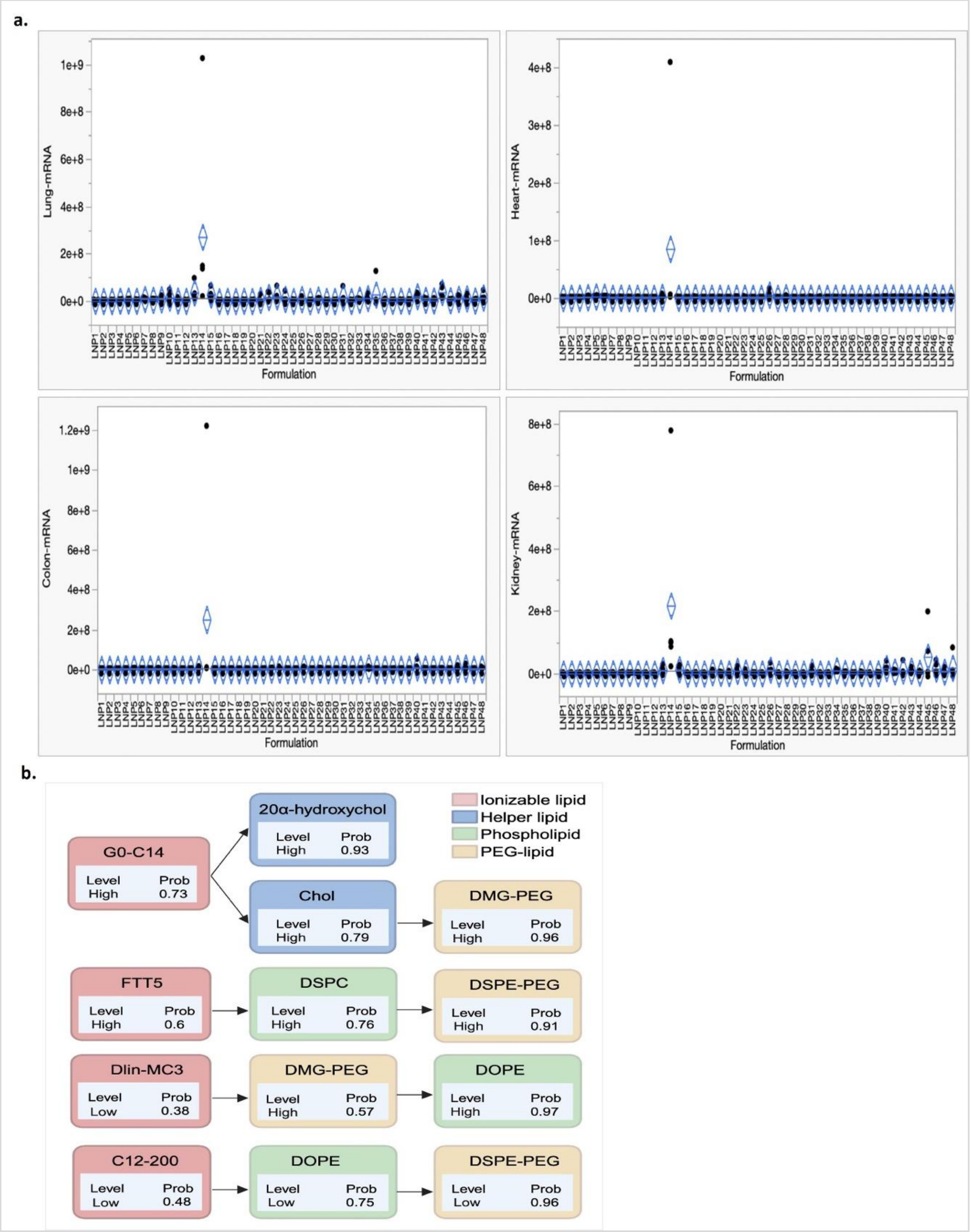


**Fig. S4. Statistical analysis of Fluc mRNA expression influenced by formulation composition across different organs.** (a) ANOVA results showing mRNA expression (Fluc bioluminescence) across 48 formulations in lung (top left), heart (top right), colon (bottom left) and kidney (bottom right). The y-axis represents biodistribution levels, while the x-axis represents the LNP formulations. Each column represents data from five mice for the same formulation, showing minimal variability within the same formulation but significant differences across different formulations. *(b)* Decision tree analysis categorized normalized Fluc bioluminescence intensity into high, medium, and low levels based on formulation compositions. Using the partition method in JMP software, the trees identified relationships between lipid components and Fluc bioluminescence intensity shows the analysis for pancreas biodistribution.

**Table S1. Compositions of LNP Formulations in the mRNA-LNP Library**

| LNP formulations | Ionizable Lipid | Helper Lipid | Phospholipid | PEG Lipid |
|---|---|---|---|---|
| LNP1 | C12-200 | Cholesterol | DOPE | DMG-PEG |
| LNP2 | C12-200 | β-sitosterol | DOPE | DMG-PEG |
| LNP3 | C12-200 | 20α-hydroxycholesterol | DOPE | DMG-PEG |
| LNP4 | C12-200 | Cholesterol | DOPE | DSPE-PEG |
| LNP5 | C12-200 | β-sitosterol | DOPE | DSPE-PEG |
| LNP6 | C12-200 | 20α-hydroxycholesterol | DOPE | DSPE-PEG |
| LNP7 | C12-200 | Cholesterol | DSPC | DMG-PEG |
| LNP8 | C12-200 | β-sitosterol | DSPC | DMG-PEG |
| LNP9 | C12-200 | 20α-hydroxycholesterol | DSPC | DMG-PEG |
| LNP10 | C12-200 | Cholesterol | DSPC | DSPE-PEG |
| LNP11 | C12-200 | β-sitosterol | DSPC | DSPE-PEG |
| LNP12 | C12-200 | 20α-hydroxycholesterol | DSPC | DSPE-PEG |
| LNP13 | Dlin-MC3 | Cholesterol | DOPE | DMG-PEG |
| LNP14 | Dlin-MC3 | β-sitosterol | DOPE | DMG-PEG |
| LNP15 | Dlin-MC3 | 20α-hydroxycholesterol | DOPE | DMG-PEG |
| LNP16 | Dlin-MC3 | Cholesterol | DOPE | DSPE-PEG |
| LNP17 | Dlin-MC3 | β-sitosterol | DOPE | DSPE-PEG |
| LNP18 | Dlin-MC3 | 20α-hydroxycholesterol | DOPE | DSPE-PEG |
| LNP19 | Dlin-MC3 | Cholesterol | DSPC | DMG-PEG |
| LNP20 | Dlin-MC3 | β-sitosterol | DSPC | DMG-PEG |
| LNP21 | Dlin-MC3 | 20α-hydroxycholesterol | DSPC | DMG-PEG |
| LNP22 | Dlin-MC3 | Cholesterol | DSPC | DSPE-PEG |
| LNP23 | Dlin-MC3 | β-sitosterol | DSPC | DSPE-PEG |
| LNP24 | Dlin-MC3 | 20α-hydroxycholesterol | DSPC | DSPE-PEG |
| LNP25 | FTT5 | Cholesterol | DOPE | DMG-PEG |
| LNP26 | FTT5 | β-sitosterol | DOPE | DMG-PEG |
| LNP27 | FTT5 | 20α-hydroxycholesterol | DOPE | DMG-PEG |
| LNP28 | FTT5 | Cholesterol | DOPE | DSPE-PEG |
| LNP29 | FTT5 | β-sitosterol | DOPE | DSPE-PEG |
| LNP30 | FTT5 | 20α-hydroxycholesterol | DOPE | DSPE-PEG |
| LNP31 | FTT5 | Cholesterol | DSPC | DMG-PEG |
| LNP32 | FTT5 | β-sitosterol | DSPC | DMG-PEG |
| LNP33 | FTT5 | 20α-hydroxycholesterol | DSPC | DMG-PEG |
| LNP34 | FTT5 | Cholesterol | DSPC | DSPE-PEG |
| LNP35 | FTT5 | β-sitosterol | DSPC | DSPE-PEG |
| LNP36 | FTT5 | 20α-hydroxycholesterol | DSPC | DSPE-PEG |
| LNP37 | G0-C14 | Cholesterol | DOPE | DMG-PEG |
| LNP38 | G0-C14 | β-sitosterol | DOPE | DMG-PEG |
| LNP39 | G0-C14 | 20α-hydroxycholesterol | DOPE | DMG-PEG |
| LNP40 | G0-C14 | Cholesterol | DOPE | DSPE-PEG |
| LNP41 | G0-C14 | β-sitosterol | DOPE | DSPE-PEG |
| LNP42 | G0-C14 | 20α-hydroxycholesterol | DOPE | DSPE-PEG |
| LNP43 | G0-C14 | Cholesterol | DSPC | DMG-PEG |
| LNP44 | G0-C14 | β-sitosterol | DSPC | DMG-PEG |
| LNP45 | G0-C14 | 20α-hydroxycholesterol | DSPC | DMG-PEG |
| LNP46 | G0-C14 | Cholesterol | DSPC | DSPE-PEG |
| LNP47 | G0-C14 | β-sitosterol | DSPC | DSPE-PEG |
| LNP48 | G0-C14 | 20α-hydroxycholesterol | DSPC | DSPE-PEG |

**Table S2. LNP biodistribution influenced by formulation composition across different organs.**

| Organ | Lipid Type | p-value | Significance | Most Significant Lipid | Least Significant Lipid |
|---|---|---|---|---|---|
| Pancreas | Ionizable Lipid | 0.0054 | Significant | Dlin-MC3, G0-C14 | C12-200 |
| Pancreas | Helper Lipid | 0.1103 | Not Significant | N/A | N/A |
| Pancreas | Phospho Lipid | 0.5994 | Not Significant | N/A | N/A |
| Pancreas | PEG Lipid | 0.0099 | Significant | DMG-PEG | DSPE-PEG |
| Tumor | Ionizable Lipid | 0.056 | Not Significant | N/A | N/A |
| Tumor | Helper Lipid | 0.9046 | Not Significant | N/A | N/A |
| Tumor | Phospho Lipid | <0.0001 | Significant | DSPC | DOPE |
| Tumor | PEG Lipid | <0.0001 | Significant | DSPE-PEG | DMG-PEG |
| Spleen | Ionizable Lipid | 0.0035 | Significant | FTT5 | Dlin-MC3 |
| Spleen | Helper Lipid | 0.7668 | Not Significant | N/A | N/A |
| Spleen | Phospho Lipid | 0.9392 | Not Significant | N/A | N/A |
| Spleen | PEG Lipid | 0.6648 | Not Significant | N/A | N/A |
| Liver | Ionizable Lipid | 0.0211 | Significant | FTT5 | Dlin-MC3 |
| Liver | Helper Lipid | 0.7989 | Not Significant | N/A | N/A |
| Liver | Phospho Lipid | 0.3688 | Not Significant | N/A | N/A |
| Liver | PEG Lipid | 0.2393 | Not Significant | N/A | N/A |

**Table S3. mRNA expression influenced by formulation composition across different organs.**

| Organ | Lipid Type | P Value | Significance | Most Significant Lipid | Least Significant Lipid |
|---|---|---|---|---|---|
| Pancreas | Ionizable Lipid | <0.0001 | Significant | G0-C14 | C12-200 |
| Pancreas | Phospholipid | 0.0074 | Significant | DSPC | DOPE |
| Pancreas | PEG Lipid | 0.0125 | Significant | DMG-PEG | DSPE-PEG |
| Pancreas | Helper Lipid | 0.0710 | Significant | Cholesterol | 20α-hydroxycholesterol |
| Tumor | Ionizable Lipid | <0.0001 | Significant | G0-C14 | C12-200 |
| Tumor | Phospholipid | <0.0001 | Significant | DSPC | DOPE |
| Tumor | PEG Lipid | 0.0298 | Significant | DSPE-PEG | DMG-PEG |
| Tumor | Helper Lipid | 0.9305 | Not Significant | N/A | N/A |
| Spleen | Ionizable Lipid | 0.6448 | Not Significant | N/A | N/A |
| Spleen | PEG Lipid | 0.2037 | Not Significant | N/A | N/A |
| Spleen | Phospholipid | 0.0032 | Significant | DSPC | DOPE |
| Spleen | Helper Lipid | 0.0428 | Significant | Cholesterol | Beta-Sitosterol |
| Liver | Ionizable Lipid | <0.0001 | Significant | C12-200 | G0-C14 |
| Liver | Phospholipid | <0.0001 | Significant | DOPE | DSPC |
| Liver | PEG Lipid | 0.0004 | Significant | DSPE-PEG | DMG-PEG |
| Liver | Helper Lipid | 0.0514 | Not Significant | N/A | N/A |

**Table S4. Correlation analysis of mRNA expression and RhB intensities across various organs**

| Organ | Correlation | Significance |
|---|---|---|
| Pancreas RhB – Tumor RhB | 0.1718 | Not significant |
| Pancreas RhB – Liver RhB | -0.4024 | Moderate negative correlation |
| Liver RhB – Tumor RhB | -0.1311 | Not significant |

| Organ | Correlation | Significance |
|---|---|---|
| Pancreas mRNA – Tumor mRNA | 0.2790 | Weak positive correlation |
| Pancreas mRNA – Liver mRNA | -0.6896 | Strong negative correlation |
| Liver mRNA – Tumor mRNA | -0.4204 | Moderate negative correlation |

| Organ | Correlation | Significance |
|---|---|---|
| Pancreas RhB – Pancreas mRNA | 0.3305 | Moderate positive correlation |
| Tumor RhB – Tumor mRNA | 0.4798 | Moderate positive correlation |
| Liver RhB – Liver mRNA | 0.2534 | Weak positive correlation |

**Table S5. LNP biodistribution and mRNA expression influenced by LNP size and zeta**

Table: LNP size to LNP biodistribution

| Organ | Correlation | Significance |
|---|---|---|
| Pancreas | -0.0928 | Not significant |
| Tumor | 0.2467 | Weak positive correlation |
| Liver | -0.0962 | Not significant |

Table: LNP size to mRNA expression

| Organ | Correlation | Significance |
|---|---|---|
| Pancreas | 0.0862 | Not significant |
| Tumor | 0.2816 | Weak positive correlation |
| Liver | -0.1763 | Weak negative correlation |

Table: Zeta potential to LNP biodistribution

| Organ | Correlation | Significance |
|---|---|---|
| Pancreas | 0.0338 | Not significant |
| Tumor | 0.0721 | Not significant |
| Liver | -0.0290 | Not significant |

Table: Zeta potential to mRNA expression

| Organ | Correlation | Significance |
|---|---|---|
| Pancreas | 0.0951 | Not significant |
| Tumor | 0.2103 | Weak positive correlation |
| Liver | 0.0051 | Not significant |

**Table S6. Histological analysis of liver and lung tissues for *in vivo* toxicity profiling**

| Treatment groups | Animal no. | Tissue | Observations | Tissue | Observations |
|---|---|---|---|---|---|
| Control | 1 | Liver | Tumor in liver and on capsule | Lung | Normal |
| Control | 2 | Liver | Small tumor on capsule | Lung | Normal |
| LNP7 | 1 | Liver | Tumor in adjacent fat | Lung | Dilated bronchioles |
| LNP7 | 2 | Liver | Normal | Lung | Focal increased alveolar macrophages |
| LNP7 | 3 | Liver | Tumor in adjacent fat | Lung | Normal |
| LNP14 | 1 | Liver | Tumor in liver and on capsule | Lung | Normal |
| LNP14 | 2 | Liver | Normal | Lung | Focal increase alveolar macrophages |
| LNP14 | 3 | Liver | Normal | Lung | Focal increased alveolar macrophages and dilated bronchioles |
| LNP25 | 1 | Liver | Tumor on capsule and in muscle | Lung | Normal |
| LNP25 | 2 | Liver | Necrosis; increased apoptotic cells | Lung | Focal increase alveolar macrophages |
| LNP25 | 3 | Liver | Small focus of tumor in adjacent fat | Lung | Focal increased alveolar macrophages |
| LNP40 | 1 | Liver | Normal | Lung | Focal increase alveolar macrophages |
| LNP40 | 2 | Liver | Small tumor focus on capsule | Lung | Normal |
| LNP40 | 3 | Liver | Normal | Lung | Focal increase alveolar macrophages |

**Table S7. LNP biodistribution and mRNA expression influenced by PDI**

| Table: PDI to LNP biodistribution | | |
|---|---|---|
| **Organ** | **Correlation** | **Significance** |
| Pancreas | -0.1278 | Not significant |
| Tumor | 0.1150 | Not significant |
| Liver | 0.0386 | Not significant |

| Table: PDI to mRNA expression | | |
|---|---|---|
| **Organ** | **Correlation** | **Significance** |
| Pancreas | 0.0667 | Not significant |
| Tumor | -0.1390 | Not significant |
| Liver | 0.0202 | Not significant |
